# Primate-expressed EPIREGULIN promotes basal progenitor proliferation in the developing neocortex

**DOI:** 10.1101/2023.08.23.554446

**Authors:** Paula Cubillos, Nora Ditzer, Annika Kolodziejczyk, Gustav Schwenk, Janine Hoffmann, Theresa M. Schütze, Razvan P. Derihaci, Cahit Birdir, Johannes E. M. Köllner, Andreas Petzold, Mihail Sarov, Ulrich Martin, Katherine R. Long, Pauline Wimberger, Mareike Albert

**Affiliations:** Center for Regenerative Therapies Dresden, TUD Dresden University of Technology, 01307 Dresden, Germany; Department of Gynecology and Obstetrics, TUD Dresden University of Technology, 01307 Dresden, Germany; National Center for Tumor Diseases, Dresden, Germany; Center for feto/neonatal Health, TUD Dresden University of Technology, Dresden; Max Planck Institute of Molecular Cell Biology and Genetics, Dresden, Germany; DRESDEN-concept Genome Center, Center for Molecular and Cellular Bioengineering, TUD Dresden University of Technology, Dresden, Germany; Leibniz Research Laboratories for Biotechnology and Artificial Organs, Department of Cardiothoracic, Transplantation and Vascular Surgery, Hannover Medical School, 30625 Hannover, Germany; REBIRTH-Cluster of Excellence, Germany; Centre for Developmental Neurobiology, Institute of Psychiatry, Psychology and Neuroscience, King’s College London, London SE1 1UL, United Kingdom; MRC Centre for Neurodevelopmental Disorders, King’s College London, London SE1 1UL, United Kingdom

## Abstract

Neocortex expansion during evolution is linked to higher numbers of neurons thought to result from increased proliferative capacity and neurogenic potential of basal progenitor cells (BPs) during development. Here we show that *EREG*, encoding the growth factor EPIREGULIN, is expressed in the human developing neocortex and in gorilla organoids, but not in the mouse neocortex. Addition of EPIREGULIN to the mouse neocortex increases proliferation of BPs via EGFR-mediated signaling, whereas ablation of *EREG* in human cortical organoids reduces BP proliferation. Addition of EPIREGULIN to cortical organoids promotes a further increase in proliferation of gorilla but not human BPs. Finally, we identify putative cis-regulatory elements that may contribute to inter-species differences in *EREG* expression. Overall, our results suggest that species-specific expression of EPIREGULIN may contribute to increased neocortex size in primates by providing a pro-proliferative signal to BPs in the subventricular zone progenitor niche.

## INTRODUCTION

The neocortex is central to higher cognitive functions. Mammalian brain evolution is associated with an increase in the expansion and folding of the neocortex (Llinares-Benadero and Borrell, 2019; Rakic, 2009; Sousa et al., 2017), yet our understanding of the exact mechanism of this evolutionary expansion and the genomic basis of neocortex evolution is still limited. Inter-species differences in neocortex size are mostly related to the number of neurons, which are generated from neural stem and progenitor cells (NPCs) during development. While the process of neurogenesis is largely conserved across mammals, differences in the proliferative capacity of NPCs, the abundance of distinct NPC types and the length of the neurogenic period are thought to be key determinants of neuron number (Dehay et al., 2015; Florio and Huttner, 2014; Lui et al., 2011; Taverna et al., 2014; Zhou et al., 2023).

NPCs can be divided into two principal groups: Apical progenitors (APs), mainly apical radial glia (aRG), that reside in the ventricular zone (VZ), and BPs that reside in the subventricular zone (SVZ). Across mammals, aRG are highly proliferative and can generate more aRG via symmetric proliferative divisions, or BPs and neurons, mostly via multiple rounds of asymmetric self-renewing divisions (Götz and Huttner, 2005; Rakic, 2003). Their position adjacent to the ventricle provides aRG access to the pro-proliferative signals of the cerebrospinal fluid (Lehtinen et al., 2011).

In contrast, BPs are variable in their proliferative potential across mammals. In species with a small and smooth neocortex, such as the mouse, BPs typically divide only once to generate two neurons (Haubensak et al., 2004; Miyata et al., 2004; Noctor et al., 2004), whereas BPs of species with a large and folded neocortex, such as human, can undergo multiple rounds of divisions (Betizeau et al., 2013; Florio and Huttner, 2014; Lui et al., 2011). Species differences are also linked to the types of BPs that are present. In the mouse neocortex (mNcx), basal intermediate progenitor cells (bIPs) are predominant, whereas in species with a large neocortex, basal or outer radial glia (bRG/oRG) are highly abundant (Fietz et al., 2010; Hansen et al., 2010; Reillo et al., 2011). In the context of neocortex expansion, bRG are thought to be important as they are highly proliferative (Betizeau et al., 2013; Borrell and Calegari, 2014; Florio and Huttner, 2014). Overall, the increased proliferative capacity of BPs in species with a large and folded neocortex leads to an expansion of the SVZ, resulting in an inner and outer layer (ISVZ/OSVZ), which represents an additional proliferative niche away from the ventricle and the signals of the cerebrospinal fluid (Fietz et al., 2012; Florio and Huttner, 2014; Lehtinen et al., 2011; Libe-Philippot and Vanderhaeghen, 2021; Pollen et al., 2015). While extracellular matrix (ECM) components are thought to contribute to this niche (Kalebic and Huttner, 2020), few other factors have been functionally explored.

Recent research addressing the genomic basis of human, or generally primate, neocortex expansion has led to the identification of a limited number of human- and primate-specific genes that are implicated in NPC proliferation and/or neocortex expansion and folding (Fiddes et al., 2018; Florio et al., 2015; Florio et al., 2018; Ju et al., 2016; Liu et al., 2017; Pinson et al., 2022; Suzuki et al., 2018; Van Heurck et al., 2022; Zhou et al., 2023). In addition to such novel gene functions that mostly arose from recent segmental duplication events (Dennis and Eichler, 2016), phenotypic evolution is thought to be driven by changes in developmental gene regulatory networks (Davidson and Erwin, 2006). Comparative transcriptomic studies of mouse and primates have provided evidence of primate- or human-specific gene expression changes during cortical development, suggesting that non-coding regulatory modifications contribute to brain evolution (Doan et al., 2018; Libe-Philippot and Vanderhaeghen, 2021; Mitchell and Silver, 2018). Specifically, large deletions and duplications in non-coding regions (McLean et al., 2011), human accelerated regions (HARs) (Hubisz and Pollard, 2014; Whalen et al., 2023) and human ancestor quickly evolved regions (HAQERs) (Mangan et al., 2022) have been reported to involve neurodevelopmental enhancers. *HARE5* is an exciting example of an enhancer with species-specific activity in the developing neocortex, linked to *FZD8* encoding a receptor in the canonical Wnt signaling pathway (Boyd et al., 2015). Despite the relevance of such inter-species differences in gene expression arising from changes in gene regulatory networks, few other primate- or human-expressed genes have been functionally explored to date (Libe-Philippot and Vanderhaeghen, 2021; Mitchell and Silver, 2018).

Here we study EPIREGULIN, a member of the epidermal growth factor (EGF) family of ligands that drive multiple cellular signal transduction pathways, such as ERK and AKT activation, through the ErbB subclass of receptor tyrosine kinases (Abud et al., 2021; Hynes and Lane, 2005). EPIREGULIN is encoded by the *EREG* gene and is first synthesized as a membrane-anchored precursor that is then released from the cell surface by the metalloproteinase ADAM17 (Sahin et al., 2004).

Although *Ereg* knockout mice do not show any overt developmental phenotypes (Lee et al., 2004), Epiregulin has been implicated in the regulation of cell proliferation, differentiation and cell death in multiple contexts, such as angiogenesis, skin inflammation, ovarian follicle formation and cancer (Riese and Cullum, 2014). In this study, we have examined the role of EPIREGULIN in NPC proliferation in the context of neocortex evolution by manipulating EPIREGULIN levels in the mNcx, gorilla cortical organoids and the developing human neocortex (hNcx). We propose that EPIREGULIN secretion by radial glia of the developing hNcx contributes a pro-proliferative signal to the niche of the SVZ that contributes to basal progenitor amplification.

## RESULTS

### *EREG* is expressed in human but not mouse radial glia of the developing neocortex

We previously performed inter-species transcriptome comparisons of sorted NPCs from the developing mNcx and hNcx, resulting in the identification of 207 genes that are expressed in human radial glia (RG) at higher levels than in neurons and that are not expressed in mouse NPCs, despite the presence of an ortholog in the mouse genome (Florio et al., 2015). Of these, 62 genes were marked by repressive histone 3 lysine 27 tri-methylation (H3K27me3) in the mNcx (Albert et al., 2017), potentially contributing to their repression in the mouse.

Among these genes was *EREG*, encoding for the growth factor EPIREGULIN (Figure 1A), which was undetectable by RNA-seq in the mNcx (Figure 1B), but expressed in human aRG and bRG at higher levels than in neurons in fetal hNcx at 12/13 and 18/19 gestation weeks (GW) (Figure 1C, S1A) (Albert et al., 2017; Florio et al., 2015; Johnson et al., 2015). Moreover, *EREG* was also expressed in aRG of human cerebral organoids (Figure S1B) (Camp et al., 2015). The absence of *Ereg* mRNA expression in the mNcx was supported by *in situ* hybridization data (Figure S1C) (Allen Institute for Brain Science, 2004). Expression analysis in embryonic mNcx and fetal hNcx tissue by RT-qPCR (Figure S1D, E) corroborated the inter-species difference in *EREG* mRNA expression.

**Figure 1.**
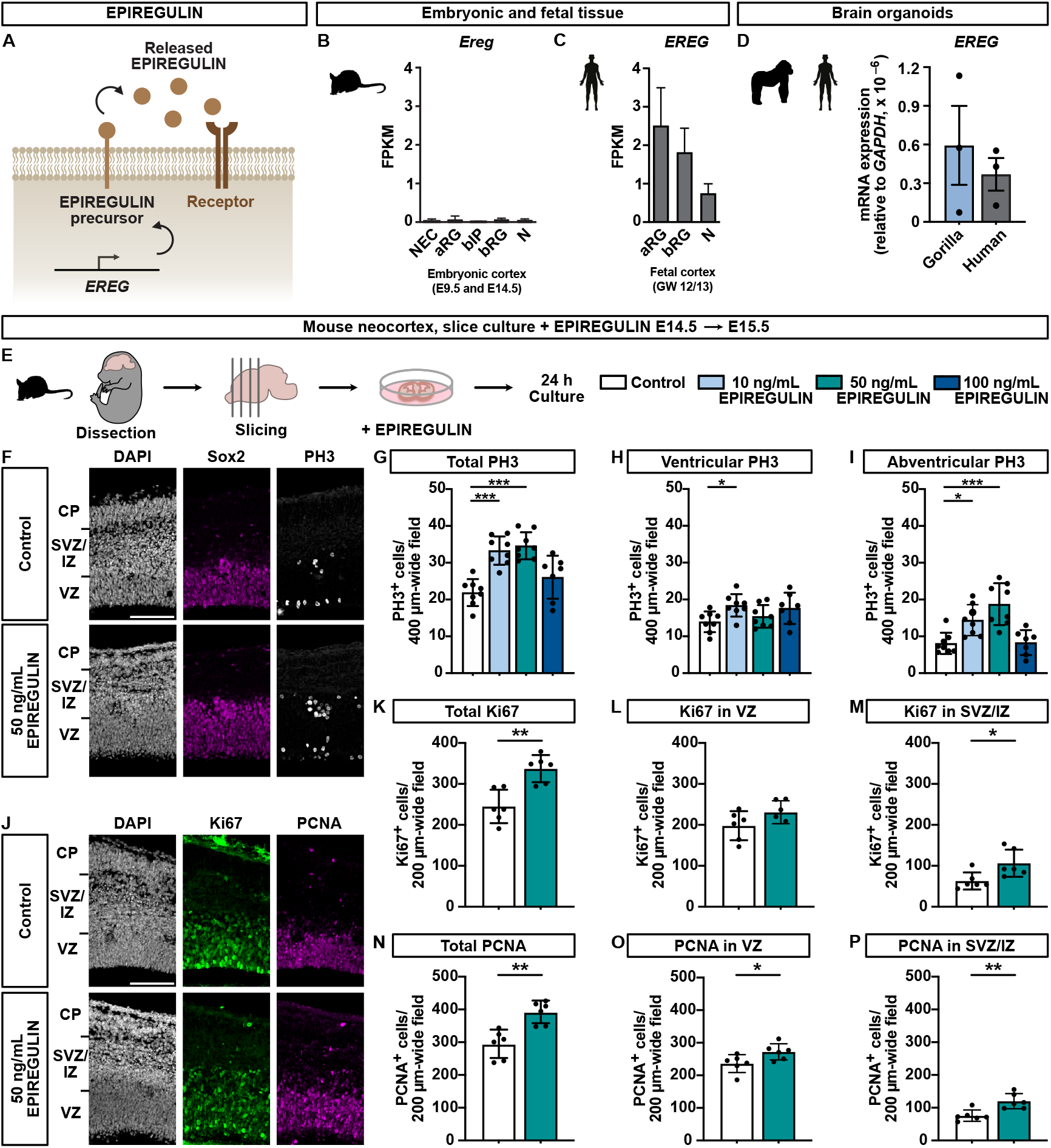
*EREG* is expressed in the developing hNcx and induces NPC proliferation in the mNcx. (A) Schematic illustration of the expression of EPIREGULIN from the *EREG* gene. (B, C) *EREG* mRNA levels in mouse (B) and human (C) NPCs and neurons analyzed by RNA-seq (data from Albert et al. (2017); Florio et al. (2015)). (D) RT-qPCR expression analysis of *EREG* in gorilla cerebral organoids (week 8) and human cortical organoids (week 6), relative to *GAPDH*. Error bars represent SD of three organoids. (E) Schematic of experimental workflow. Mouse brains (E14.5) were cut into slices and treated with different concentrations of EPIREGULIN for 24 h. (F) DAPI staining and immunofluorescence for Sox2 and PH3 of slices treated with 50 ng/mL EPIREGULIN. (G–I) Quantifications of total (G), ventricular (H) and abventricular (I) mitotic PH3-positive cells. (J) DAPI staining and immunofluorescence for Ki67 and PCNA of slices treated with 50 ng/mL EPIREGULIN. (K–M) Quantifications of total Ki67-positive cells (K), and Ki67-positive cells in the VZ (L) and SVZ/IZ (M). (N–P) Quantifications of total PCNA-positive cells (N), and PCNA-positive cells in the VZ (O) and SVZ/IZ (P). Scale bars, 100 μm. Error bars represent SD; *** p < 0.001, ** p < 0.01, * p < 0.05; one-way ANOVA with Dunnett post hoc test (G–I), and Student’s *t*-test (K–P).

Furthermore, in the mNcx, the *Ereg* gene locus showed low enrichment for active chromatin modifications, such as acetylation of H3K27 (H3K27ac) and H3K4me3, and instead was enriched in repressive H3K27me3 (Figure S1F). In contrast, the human *EREG* locus was highly enriched in active H3K27ac in the fetal hNcx, in agreement with the differential *EREG* expression between the mNcx and hNcx.

### *EREG* is expressed in primate neural progenitor cells

To address whether expression of *EREG* in the developing neocortex is a human-specific feature or is also found in other primates, we generated gorilla induced pluripotent stem cell (iPSC)-derived cerebral organoids and compared *EREG* expression to human iPSC-derived organoids (Figure 1D, S1G). Both gorilla and human organoids expressed *EREG* mRNA at similar levels during neurogenesis. Moreover, mining of RNA-seq data from macaque and human NPCs (Kliesmete et al., 2023) revealed similar expression levels of *EREG* mRNA in both species (Figure S1H). These data indicate that *EREG* expression is not restricted to humans but also seen in two additional gyrencephalic primates.

Given the differential expression of *EREG* in the developing neocortex of a species with a small and smooth neocortex versus the expanded and highly folded primate neocortex, we speculated that the growth factor EPIREGULIN might contribute to differences in the proliferative capacity of NPCs between species.

### Addition of EPIREGULIN to the developing mNcx increases basal progenitor proliferation

Evidence for a potential role of *EREG* in cell proliferation comes from data of human brain tumors (Figure S2). While *EREG* expression is not *per se* increased in glioblastoma, *EREG* expression is higher in high grade compared to low grade glioma and loss of *EREG* correlates with increased survival, supporting a potential role of EPIREGULIN in cell proliferation. Furthermore, Epiregulin was shown to enhance tumorigenicity by activating the ERK/MAPK pathway in a mouse model of glioblastoma (Kohsaka et al., 2014).

To directly test the effect of EPIREGULIN on NPC proliferation, we incubated organotypic slice cultures of embryonic day (E) 14.5 mNcx with or without different concentrations of recombinant EPIREGULIN for 24 hours (Figure 1E), followed by analysis of cell proliferation. Immunofluorescence of phospho-histone 3 (PH3), a marker of mitotic cells, revealed that 10 ng/mL and 50 ng/mL of EPIREGULIN caused a significant increase in the total number of mitotic cells (Figure 1F, G). While there was only a small increase in ventricular mitosis (Figure 1H), abventricular mitosis was strongly increased (Figure 1I), suggesting that BPs are particularly responsive to the addition of EPIREGULIN. Of the three EPIREGULIN concentrations tested, the intermediate concentration of 50 ng/mL resulted in the highest increase in abventricular PH3-positive cells compared to the control condition. This concentration was therefore used for all further experiments.

We next examined additional markers of cell proliferation, specifically Ki67 and proliferating cell nuclear antigen (PCNA) (Figure 1J–P). Total numbers of both Ki67 and PCNA expressing cells were increased upon addition of EPIREGULIN (Figure 1K, N). While there was only a small increase in Ki67 and PCNA in the VZ (Figure 1L, O), both were significantly increased in the SVZ/IZ (Figure 1M, P). These data suggest that EPIREGULIN stimulates proliferation of NPCs, particularly in the SVZ, at mid-neurogenesis in the mNcx.

### Addition of EPIREGULIN to the developing mNcx increases both major basal progenitor types

Next, we explored which of the distinct NPC types were affected by the addition of EPIREGULIN. First, we performed immunofluorescence analysis of the RG marker Sox2 in organotypic slice cultures with and without EPIREGULIN treatment. The total number of Sox2-positive cells was higher after addition of EPIREGULIN (Figure 2A, B), which was mainly driven by a significant increase in the number of Sox2-positive cells in the SVZ/IZ (Figure 2C, D). In addition, we observed an increase in the total number of Tbr2-positive bIPs (Figure 2E–G), particularly in the VZ, which may represent newborn bIPs.

**Figure 2.**
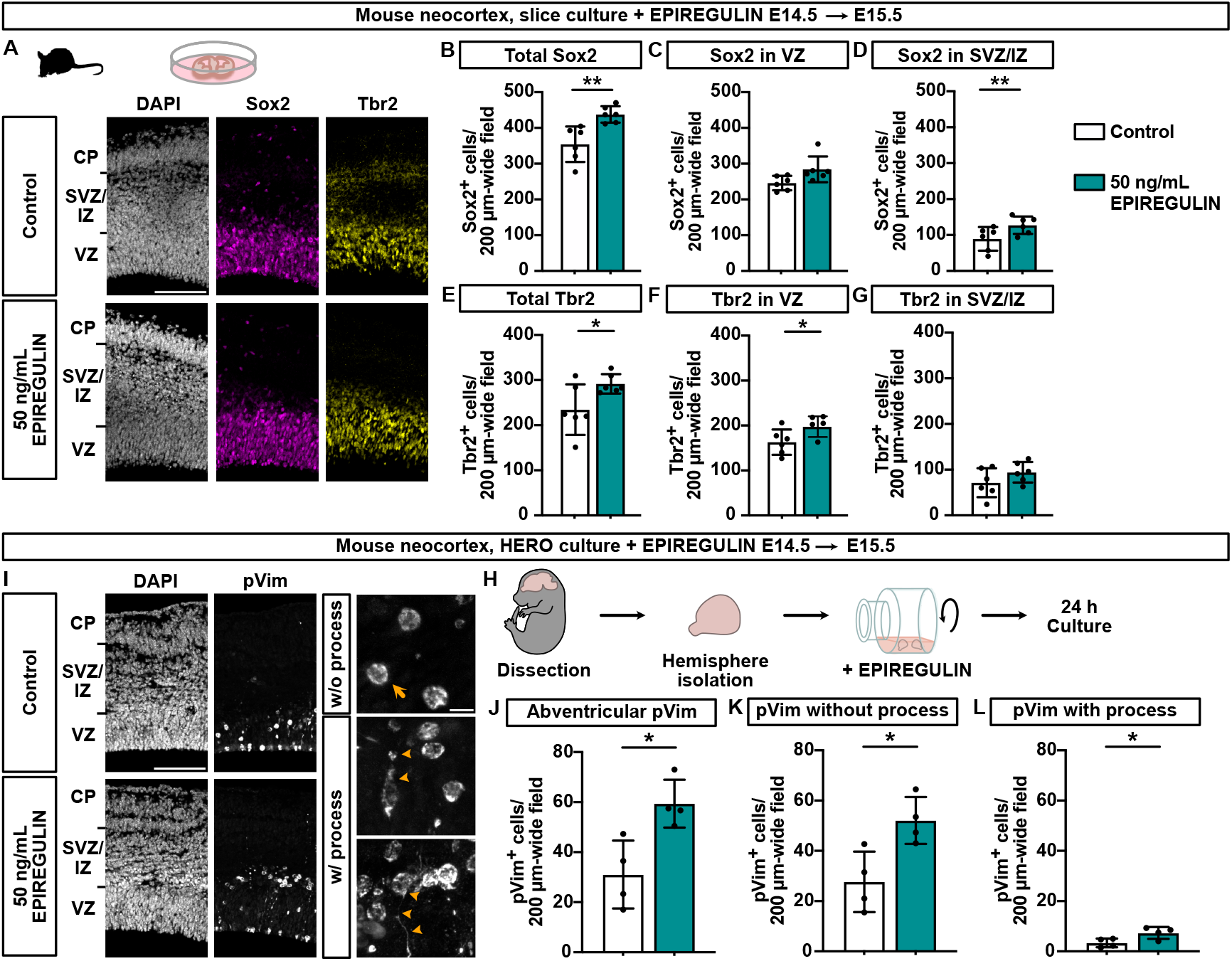
Addition of EPIREGULIN to the mNcx increases BPs. (A) DAPI staining and immunofluorescence for Sox2 and Tbr2 of mouse slices (E14.5) cultured with 50 ng/mL EPIREGULIN for 24 h. (B–D) Quantifications of total Sox2 (B), and Sox2 in the VZ (C) and SVZ/IZ (D). (E–G) Quantifications of total Tbr2 (E), and Tbr2 in the VZ (F), and SVZ/IZ (G). (H) Schematic of experimental workflow. Mouse brain hemispheres (E14.5) were cultured under rotation (HERO) in the presence of 50 ng/mL EPIREGULIN for 24 h. (I) DAPI staining and immunofluorescence for phospho-Vimentin (pVim) to distinguish mitotic BPs without (top; arrow) or with (bottom; arrowheads) a process. (J–L) Quantifications of abventricular pVim-positive cells (J), and pVim-positive cells without (K) and with (L) a process. Scale bars, 100 μm (A, I left) and 10 μm (I inset). Error bars represent SD; ** p < 0.01, * p < 0.05; Student’s *t*-test.

A high number of Sox2-positive bRG is a characteristic of the expanded SVZ in gyrencephalic species, such as human (Fietz et al., 2010; Hansen et al., 2010; Reillo et al., 2011). To further characterize the BP population in the EPIREGULIN-treated mNcx, we performed hemisphere rotation (HERO) cultures at E14.5 (Figure 2H) (Schenk et al., 2009), followed by slicing into 70 μm thick sections to facilitate the analysis of basal processes, another hallmark of bRG (Fietz et al., 2010; Hansen et al., 2010; Reillo et al., 2011). Using phospho-Vimentin (pVim) to analyze mitotic BPs, we could distinguish BPs without a process, mostly bIPs, and with a process, mostly bRG (Figure 2I). This analysis corroborated the increase in abventricular mitosis (Figure 2J) already observed with PH3 (Figure 1I), and further revealed that mitotic cells both with and without a process were increased upon EPIREGULIN treatment (Figure 2K, L).

Taken together, we observed that the addition of recombinant EPIREGULIN to mNcx cultures resulted in an increase in NPC proliferation, preferentially in the SVZ. Both major types of BPs, bIPs characterized by Tbr2 expression and the lack of a process as well as bRG marked by Sox2 and the presence of a process, were increased, suggesting that EPIREGULIN can induce the amplification of different BP types in the mNcx.

### EPIREGULIN ablation in human cortical organoids reduces basal progenitor proliferation

We next examined whether endogenously expressed EPIREGULIN is required for BP proliferation in the hNcx. To address this important point, we used sliced human cortical organoids, which were reported to sustain neurogenesis, organoid growth and SVZ expansion over a long time-period (Qian et al., 2020). We considered organoids to be a suitable model for this question as *EREG* expression was also observed in human cerebral and cortical organoids in addition to human fetal tissue (Figure 1D, S1A). We employed the CRISPR/Cas9 system and injected guide RNAs (gRNA) in complex with recombinant Cas9 protein (Kalebic et al., 2016) into ventricle-like structures of 6-week cortical organoids followed by electroporation (EP) (Figure 3A). Co-electroporation of a GFP expression plasmid allowed the identification of targeted cells (Figure 3B). To disrupt the expression of *EREG*, a set of two gRNAs was used that showed efficient targeting *in vitro* and in the iPSC line used to generate the cortical organoids (Figure S3). Seven days post EP, immunofluorescence for the proliferation marker KI67 was performed (Figure 3C–E). This showed a significant reduction in the percentage of KI67-positive GFP-positive cells in the SVZ/CP-like region (Figure 3E), but not in the VZ (Figure 3D), compared to the control. This indicates that EPIREGULIN is required for BP, but not AP, proliferation in human cortical organoids.

**Figure 3.**
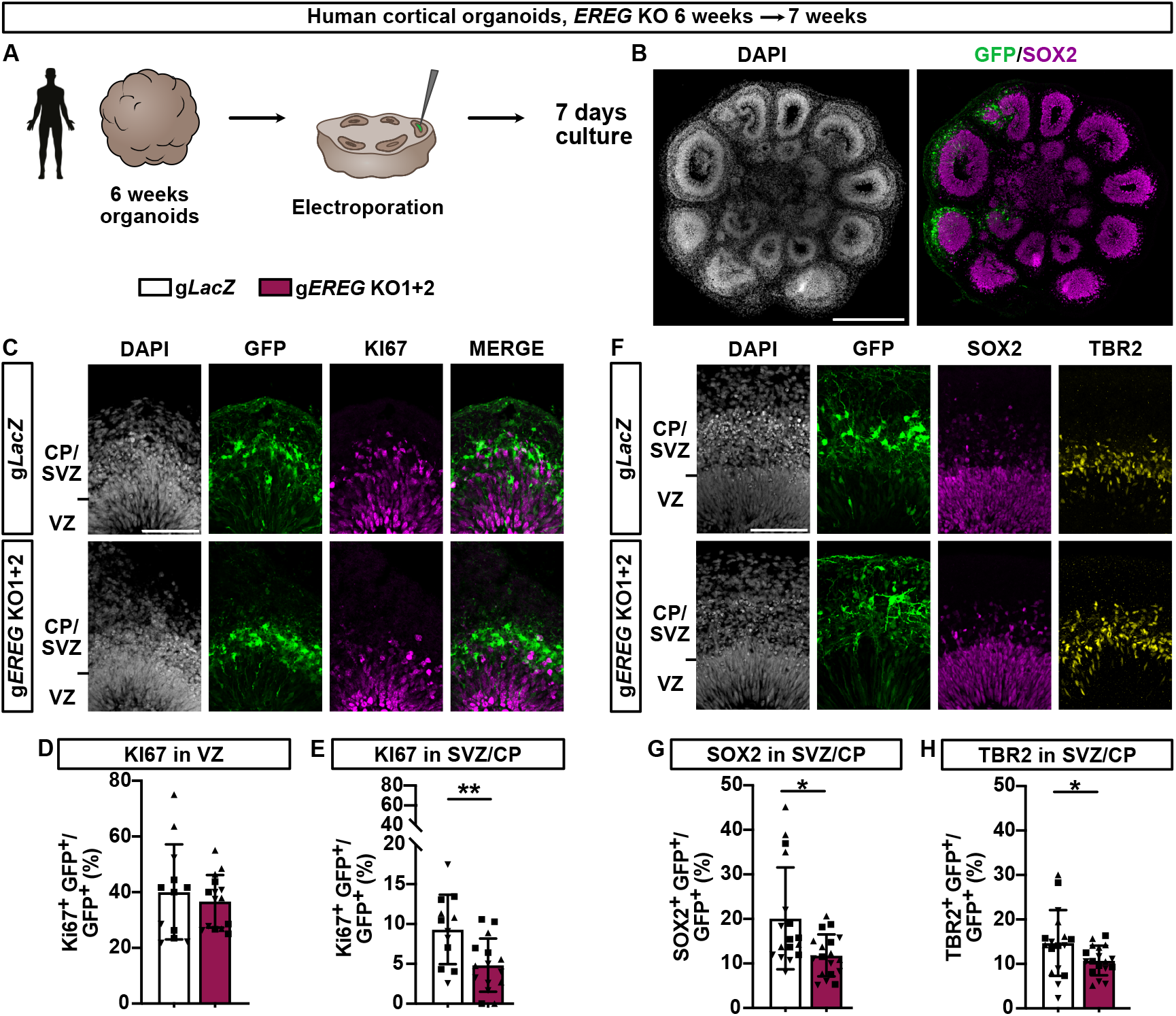
EPIREGULIN ablation in human cortical organoids reduces BP proliferation. (A) Schematic of experimental workflow. Human cortical organoids (6 weeks) were electroporated with a plasmid encoding GFP and CRISPR/Cas9 RNP complexes targeting either *LacZ* or *EREG*, and analyzed after 7 days. (B) DAPI staining and immunofluorescence for GFP and SOX2 of an electroporated human cortical organoid. (C) DAPI staining and immunofluorescence for GFP and KI67. (D–E) Quantifications of KI67 in the VZ (D) and SVZ/CP (E). (F) DAPI staining and immunofluorescence for SOX2 and TBR2. (D–E) Quantifications of SOX2 (G) and TBR2 (H) in the SVZ/CP. Scale bars, 500 μm (B) and 100 μm (C, F). Error bars represent SD; ** p < 0.01, * p < 0.05; one-way ANOVA with Dunnett post hoc test.

To assess which NPC types were affected by ablation of *EREG* in the SVZ, we performed immunofluorescence staining for the RG marker SOX2 and the bIP marker TBR2 (Figure 3F– H). Both SOX2 and TBR2 were reduced upon *EREG* targeting (Figure 3G, H), suggesting that EPIREGULIN contributes to the proliferation of both BP types, bIPs and bRG, in 6-week cortical organoids, in which the SVZ-like area is in the process of expansion.

### Addition of EPIREGULIN to gorilla, but not human, cortical organoids increases basal progenitor proliferation

Given that exogenous EPIREGULIN promoted BP proliferation in the mNcx, we next asked whether the addition of EPIREGULIN to the hNcx would increase BP proliferation even further. To address this question, we first incubated human fetal cortical tissue of 12/13 GW in a free-floating tissue culture (FFTC) system (Long et al., 2018) with and without 50 ng/mL EPIREGULIN for 24 hours (Figure 4A). Immunofluorescence analysis showed no change in the proliferation marker KI67 nor in the NPC markers SOX2 and TBR2 (Figure 4B–E). Considering that the cell cycle length of primate NPCs is much longer than in the mNcx (Kornack and Rakic, 1998), we speculated that the 24-hour incubation time might not have been sufficient to induce a measurable increase in proliferation. We therefore turned to the cortical organoid system again, which allowed addition of EPIREGULIN for longer time periods and at different developmental stages. To rule out any issues of EPIREGULIN penetration to the organoid core, we added the growth factor directly after organoid slicing when the ventricle-like structures are open and accessible. However, neither a 4- or 10-day incubation with EPIREGULIN was sufficient to increase NPC proliferation or NPC numbers in 6-week (data not shown) or 10-week human cortical organoids (Figure 4F–J).

**Figure 4.**
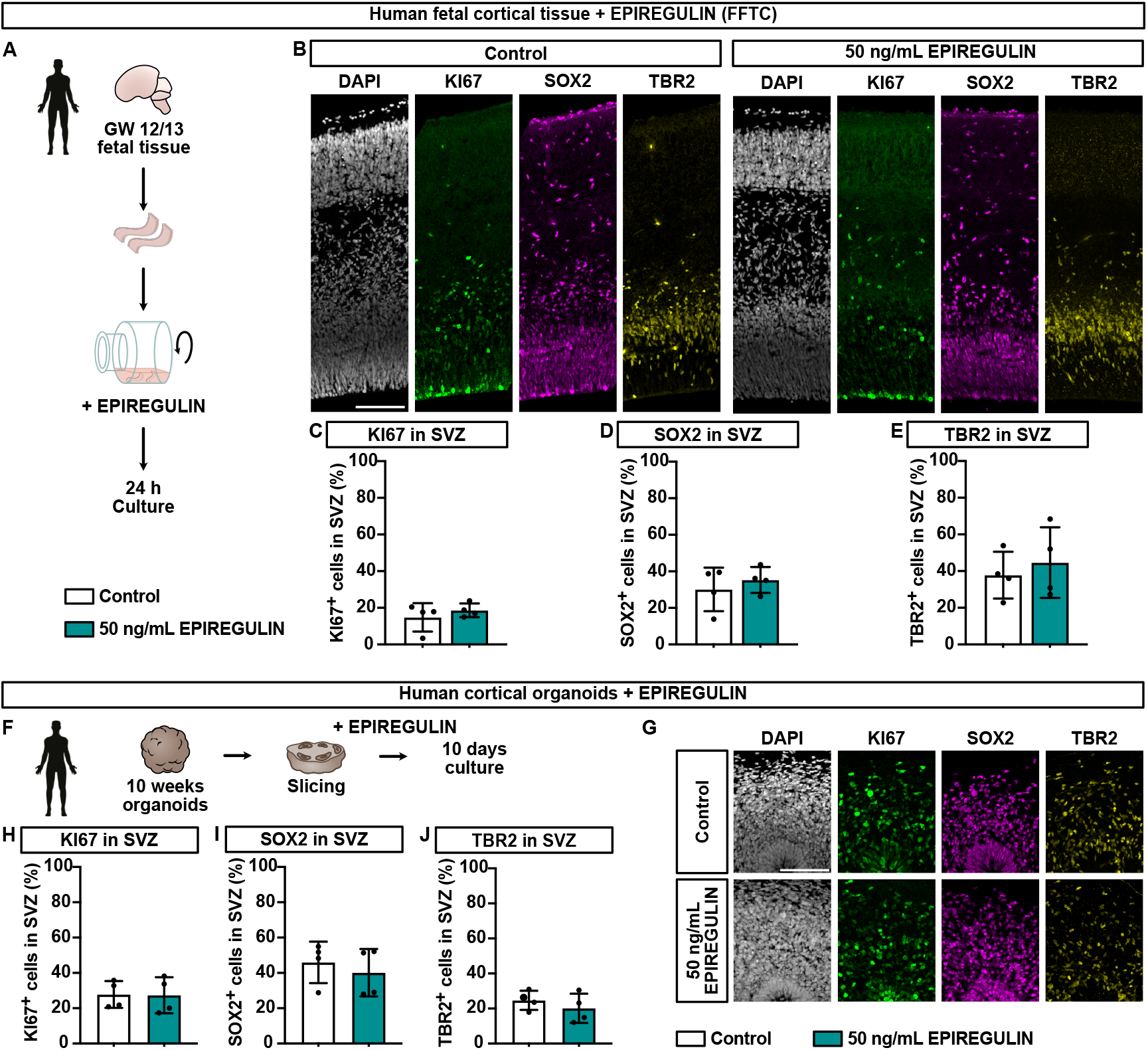
Addition of EPIREGULIN to the hNcx does not further induce proliferation. (A) Schematic of experimental workflow. Human fetal tissue pieces (GW 12/13) were isolated and cultured in a free-floating tissue culture (FFTC) in the presence of 50 ng/mL of EPIREGULIN for 24 h. (B) DAPI staining and immunofluorescence for KI67, SOX2 and TBR2. (C–E) Quantifications of KI67 (C), SOX2 (D) and TBR2 (E) in the SVZ. (F) Schematic of experimental workflow. Human sliced cortical organoids (10 weeks) were treated with 50 ng/mL of EPIREGULIN for 10 days. (G) DAPI staining and immunofluorescence for KI67, SOX2 and TBR2 of human cortical organoids treated with EPIREGULIN. (H–J) Quantifications of KI67 (H), SOX2 (I) and TBR2 (J) in the SVZ of human cortical organoids. Scale bars, 100 μm. Error bars represent SD of 4 independent fetal tissue samples (C–E) or 4 cortical organoids (H–J); Student’s *t*-test; no statistically significant changes were detected.

In contrast, incubation of 6-week gorilla cortical organoids with the same concentration of EPIREGULIN resulted in a significant increase in the percentage of KI67-positive cells in the SVZ (Figure 5A–C). In particular, the percentage of SOX2-positive BPs was increased (Figure 5D) whereas the percentage of TBR2-positive BPs remained largely unchanged (Figure 5E).

**Figure 5.**
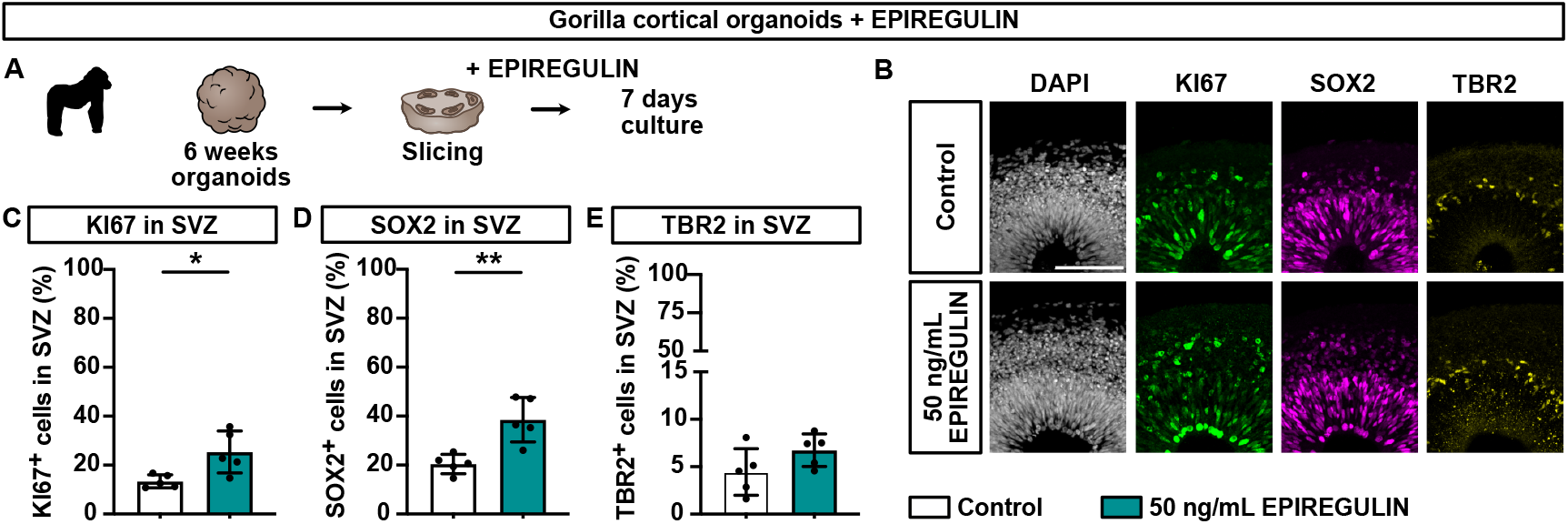
Addition of EPIREGULIN to gorilla organoids further induces BP proliferation. (A) Schematic of experimental workflow. Gorilla sliced cortical organoids (6 weeks) were treated with 50 ng/mL of EPIREGULIN for 7 days. (B) DAPI staining and IF for KI67, SOX2 and TBR2 gorilla cortical organoids treated with EPIREGULIN. (C–E) Quantifications of KI67 (C), SOX2 (D) and TBR2 (E) in the SVZ of gorilla cortical organoids. Scale bars, 100 μm. Error bars represent SD of 5 cortical organoids (C–E); ** p < 0.01, * p < 0.05; Student’s *t*-test.

Taken together, treatment with EPIREGULIN did not further increase human BP proliferation in either primary fetal tissue nor in cortical organoid cultures, suggesting that the system is already saturated and that, at least at this concentration, human BPs are not receptive to further stimulation by EPIREGULIN. Unlike the human, gorilla BP proliferation, in particular the percentage of SOX2-positive cells likely representing bRG, could be further stimulated by the addition of EPIREGULIN.

### Addition of EPIREGULIN does not induce major gene expression changes in the mNcx

To investigate the mechanism through which EPIREGULIN promotes NPC proliferation, we first analyzed gene expression changes upon addition of EPIREGULIN to the mNcx at E14.5 to identify putative down-stream target genes. Since EPIREGULIN promoted changes in the cellular composition of the mNcx (Figure 2), which would be expected to bias any gene expression analysis, we performed immuno-fluorescent activated cell sorting (FACS) to isolate neural cell populations. Specifically, RG, IPs and neurons (Figure S4A, B) were isolated based on immuno-staining for the nuclear markers Sox2 and Tbr2 (Florio et al., 2015; Schütze et al., 2023), and neurons based on expression of GFP in the *Tubb3*::GFP mouse reporter line (Attardo et al., 2008). Isolated cell populations were validated by expression of the marker genes *Sox2* and *Prom1* (Prominin 1/CD133) for aRG, *Eomes* (Tbr2) for IPs, and *Dcx* (Doublecortin) and *Tubb3* (Tuj1) for neurons (Figure S4C, D). The RNA-seq data confirmed the lack of *Ereg* expression in the mNcx (Figure S4D). Comparison of the NPC populations from control and EPIREGULIN treated mNcx did not reveal any significant gene expression changes (Figure S4E). In line with this, principal component analysis showed strong clustering of each of the three cell types, irrespective of EPIREGULIN treatment (Figure S4F). Taken together, the RNA-seq analysis of sorted NPC populations did not point to major gene expression changes as mechanism underlying the EPIREGULIN-induced increase in cell proliferation.

### EPIREGULIN competes with EGF

Secondly, we considered that secreted EPIREGULIN, released upon cleavage by ADAM17 (Figure 6A–C), might induce downstream signaling responses, via binding to its target receptors that ultimately affect the balance of NPC proliferation versus differentiation. Of note, while EPIREGULIN is one of eleven structurally related members of the EGF family of proteins (Abud et al., 2021), none of the other members showed consistent differences in gene expression between the mNcx and hNcx (Figure S5). EPIREGULIN has been reported to bind to the EGF receptor (EGFR) and ERBB4 receptor (Figure 6A, S5A) (Hynes and Lane, 2005). Mining RNA-seq data (Florio et al., 2015), we found that *EGFR* and *ERBB4* are both expressed in human aRG, bRG and neurons (Figure 6B, C). Interestingly, while *Egfr* expression is similar in the mNcx compared to the hNcx, *ErbB4* levels are very low in mouse RG, but high in bIPs and neurons. At the protein level, EGFR was reported to be expressed at high levels in both the rodent VZ and SVZ throughout cortical development (Eagleson et al., 1996).

**Figure 6.**
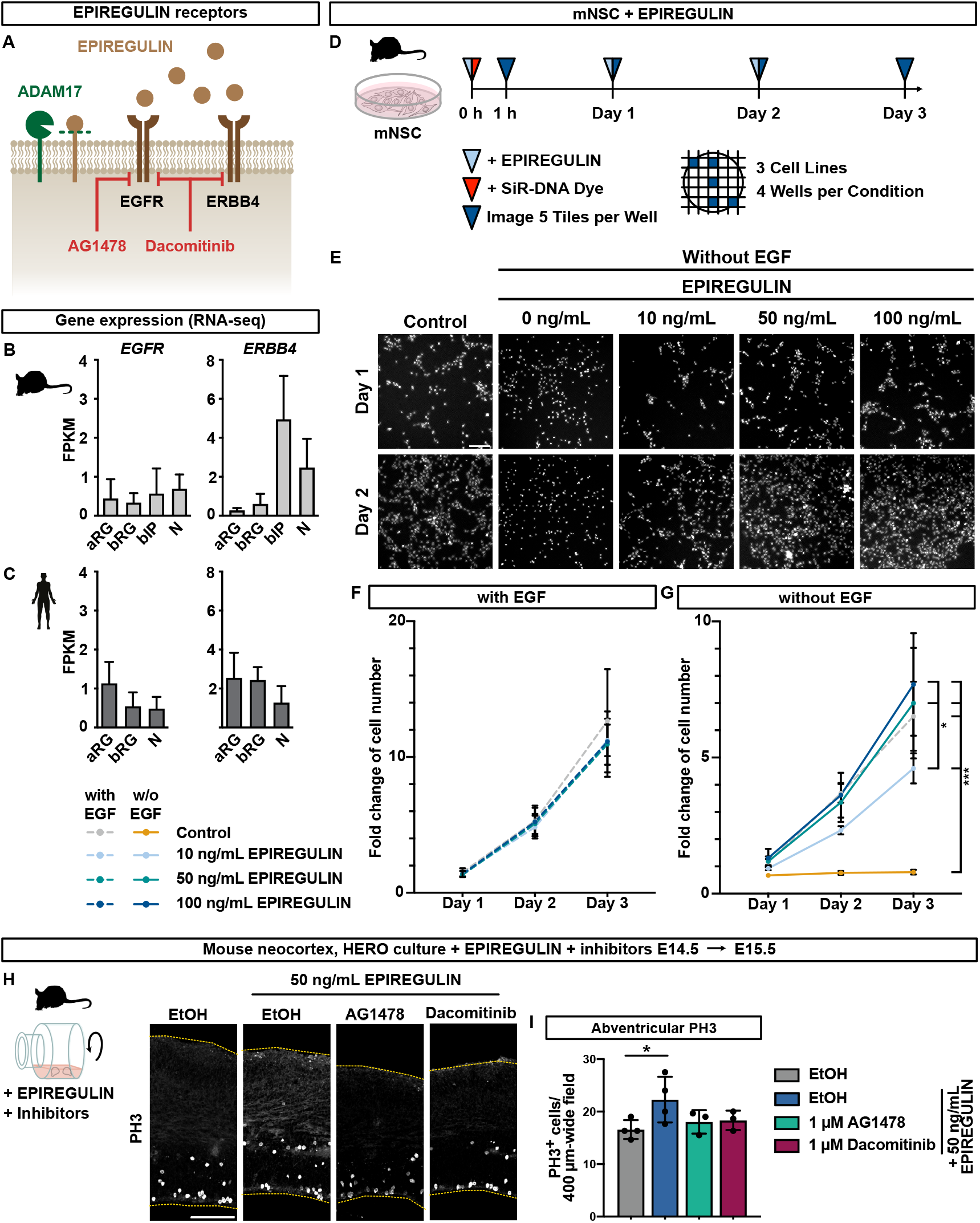
EPIREGULIN mediates BP proliferation via EGFR-signaling. (A) Schematic of the EPIREGULIN receptors and inhibitors. (B, C) Expression of EPIREGULIN receptor genes in the mNcx (B) and hNcx (C) analyzed by RNA-seq (data from Florio et al. (2015)). (D) Schematic illustration of the experimental workflow. Mouse NSC cultures were treated with different concentrations of EPIREGULIN for 3 days. The SiR-DNA dye was added to the culture to allow imaging of live cells on day 1, 2 and 3. (E) Images of SiR-DNA stained mNSC following treatment with EPIREGULIN in culture medium containing FGF but lacking EGF. The control mNSCs were cultured in medium with EGF and FGF. (F) Quantification of SiR-DNA-positive cells on day 1, 2 and 3 following EPIREGULIN treatment, shown as fold change relative to 1 hour. (G) Quantification of cells following EPIREGULIN treatment in culture medium lacking EGF. (H) Immunofluorescence for PH3 of mNcx slices treated with 50 ng/mL EPIREGULIN and receptor inhibitors for 24 hours. (I) Quantifications of abventricular mitotic PH3-positive cells. Scale bars, 100 μm. Error bars represent SD; * p < 0.05, *** p < 0.001; (F–G) two-way ANOVA; (I) one-way ANOVA with Dunnett post hoc test.

To dissect the contribution of different growth factors and receptors, we started by treating mouse neural stem cells (NSCs), derived from the dorsolateral telencephalon at E11.5/E12.5, cultured under proliferative conditions in the presence of 10 ng/mL EGF and 20 ng/mL fibroblast growth factor (FGF), with different concentrations of EPIREGULIN for three days (Figure 6D– G). This revealed that none of the different EPIREGULIN concentrations resulted in changes in mNSC proliferation *in vitro* (Figure 6F). Since EGF and EPIREGULIN are both known to bind to EGFR, we next considered that both ligands may compete for the receptor. We therefore tested the different concentrations of EPIREGULIN on mNSCs grown in medium lacking EGF (Figure 6E, G). Both EGF and FGF were reported to elicit NSC proliferation *in vitro* (Tropepe et al., 1999), and upon acute withdrawal of EGF, mNSCs did not proliferate further (Figure 6G). However, when EPIREGULIN was added, we observed a dose-dependent increase in mNSC proliferation with cells cultured with 50 ng/mL EPIREGULIN showing similar growth rates to cells grown under control conditions with 10 ng/mL EGF. Thus, EPIREGULIN can induce the proliferation of mNSC *in vitro* and this ability to stimulate proliferation is attenuated by the presence of EGF.

### EPIREGULIN promotes proliferation via the EGF receptor

Next, we aimed to investigate which receptors are required for the EPIREGULIN-induced increase in BP proliferation in the mNcx. For this, we selected two inhibitors that were reported to block signaling via the two major EPIREGULIN receptors: AG1478, an EGFR tyrosine kinase inhibitor (Martin et al., 2017) and Dacomitinib, an irreversible pan-ErbB receptor tyrosine kinase inhibitor targeting EGFR and Erbb4 (Xu et al., 2017) (Figure 6A). We then performed HERO cultures with only the solvent (ethanol; control), solvent and 50 ng/mL EPIREGULIN, or EPIREGULIN and different concentrations of the two inhibitors (Figure 6H, I). Of note, in the presence of ethanol, the EPIREGULIN-induced increase in abventricular PH3 was smaller (compare Figure 1I and 6I), but still significant. Inhibition of EGFR alone by AG1478 resulted in a complete loss of the EPIREGULIN-induced increase in abventricular mitosis (Figure 6I), whereas ventricular mitosis was surprisingly unaffected by the inhibitors (data not shown), at the concentrations used. Likewise, the pan-ErbB inhibitor Dacomitinib abrogated the increase in PH3. Taken together, these results suggest that EGFR is the key mediator of the EPIREGULIN-mediated increase in BP amplification, whereas Erbb4 appears to not play a major role. Moreover, the presence of different epidermal growth factors modulates the cellular outcome.

### Identification of putative *EREG* enhancer regions in the human genome

Finally, we aimed to dissect the mechanism of differential *EREG* expression across species. Epigenetic modifications, such as histone methylation, contribute to the regulation of gene expression during neocortex development (Bölicke and Albert, 2022; Hirabayashi and Gotoh, 2010; Hoffmann and Albert, 2021). In the mNcx, the *Ereg* locus was marked by repressive H3K27me3 and devoid of active modifications, such as H3K4me3 and H3K27ac (Figure S1F). To test whether the H3K27me3 marking was actively involved in repressing the *Ereg* gene in the mNcx, we established a CRISPR/Cas9-based epigenome editing system (Albert et al., 2017; Kearns et al., 2015) to remove H3K27me3 by targeting the catalytic jumonji domain of the histone demethylase KDMB6 (referred to as JMJC_6B) (Agger et al., 2007) to sites up- and down-stream of the *Ereg* transcription start site (TSS) (Figure S6A, B). To perform the epigenome editing, mNSCs were employed, as they recapitulate the H3K27me3 patterns observed in aRG of the mNcx (Figure S6C). Epigenome editing in the mNSCs using dCas9-JMJC_6B resulted in a reduction of H3K27me3 at the *Ereg* locus, but not at the unrelated *Hoxb5* and *Eomes* genes (Figure S6D, E). However, this loss of repressive epigenetic modifications alone did not result in an upregulation of *Ereg* gene expression (Figure S6F, G), suggesting that the mouse *Ereg* locus is in a fully repressed rather than poised state.

Given the contribution of gene regulatory networks to evolutionary changes in development (Davidson and Erwin, 2006), we next aimed to identify cis-regulatory elements (CREs) that may contribute to the expression of *EREG* in the hNcx. To this end, we examined ATAC-seq and H3K27ac ChIP-seq data of the fetal hNcx (de la Torre-Ubieta et al., 2018; Reilly et al., 2015) and found 11 regions of open chromatin within 100 kb up- and down-stream of the *EREG* gene (excluding TSS regions) that may represent putative active enhancers (Figure 7A). Two additional genes are located within this genomic region, *AREG* and *EPGN*, which also encode for growth factors known to bind to EGFR (Figure S5A). However, based on chromatin (Figure 7A) and RNA-seq (Figure S5) data, they are likely not expressed in the fetal hNcx. Using LiftOver conversion of the human genomic coordinates to the mouse genome, we identified the orthologous mouse genomic regions. Interestingly, of the 11 putative *EREG* CREs, only one shows a small ATAC-seq peak in the mNcx, whereas the other orthologous regions lack H3K27ac or ATAC-seq peaks in the mNcx (Figure 7B) (Gorkin et al., 2020). Whereas the putative CRE1 is not conserved in the mouse, the other 10 putative CREs are at least partially conserved in the mouse, yet show higher sequence divergence compared to primate species (Figure 7C).

**Figure 7.**
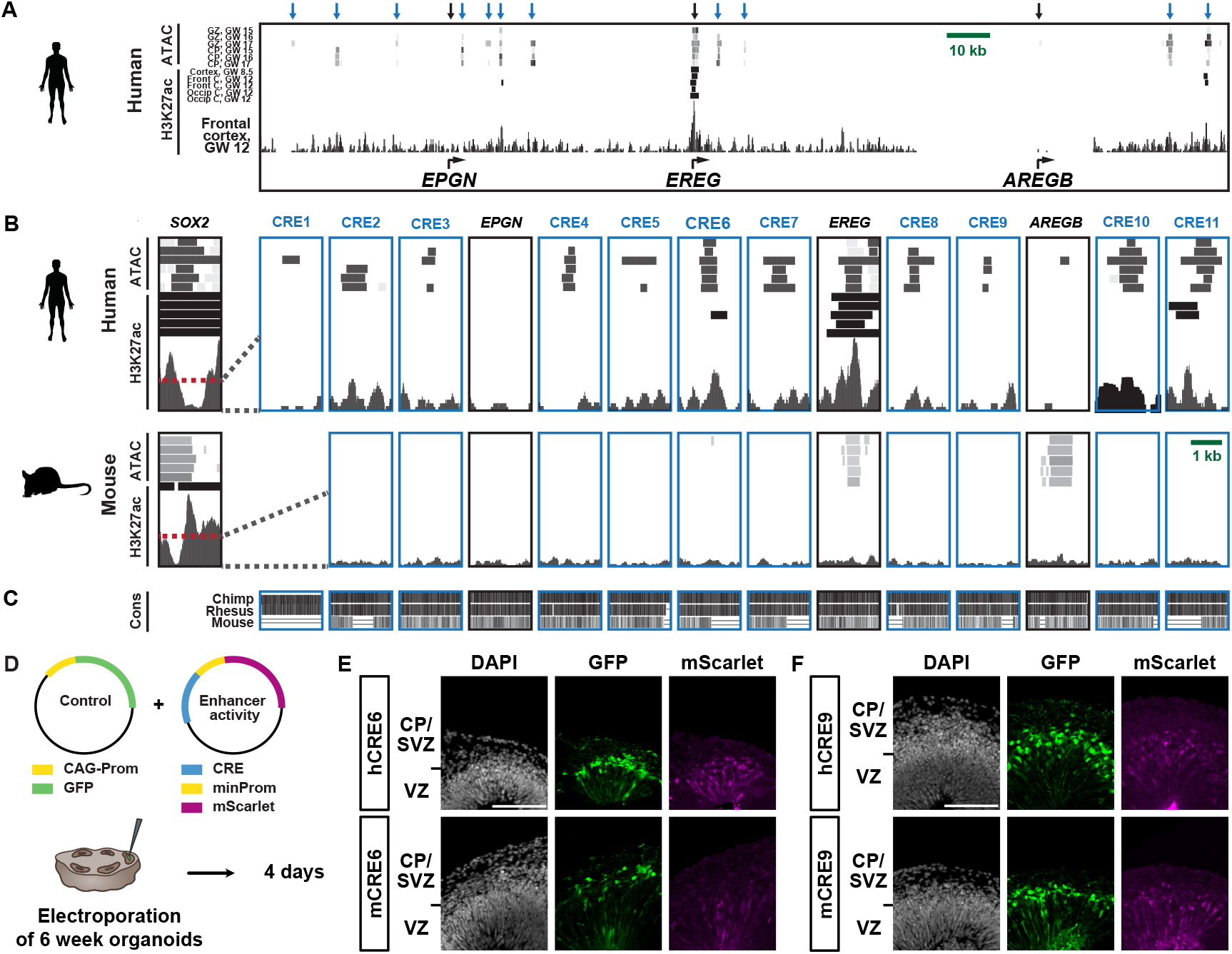
Putative enhancer regions for *EREG* differential gene expression. (A) ATAC-seq (de la Torre-Ubieta et al., 2018) and H3K27ac ChIP-seq (Reilly et al., 2015) peaks around the *EREG* gene (± 100 kb) in the hNcx at given ages. Black arrows, TSS; blue arrows, potential CREs. (B) Zoom in on the TSS and CRE regions indicated by arrows in (A), relative to open chromatin marks at the *SOX2* gene, for human (top) and mouse (bottom, (Gorkin et al., 2020)). Window size, 2 kb. (C) Evolutionary conservation from 100 vertebrate species (Blanchette et al., 2004) for the indicated species. (D) Schematic of experimental workflow. Human sliced cortical organoids (6 weeks) were electroporated with a plasmid encoding GFP and a plasmid with a putative CRE upstream of a minimal promotor driving mScarlet expression, and analyzed after 4 days. (E, F) DAPI staining and immunofluorescence for GFP and mScarlet of an electroporated human cortical organoid to test for enhancer activity of mouse and human CRE6 (E) and CRE9 (F).

To test whether these putative enhancer regions display enhancer activity, we cloned two of the human CRE regions (216 bp) and their orthologous mouse sequences, respectively, into a plasmid containing a minimal promotor upstream of a fluorescent reporter (mScarlet). These plasmids were then electroporated into human cortical organoids, along with a control plasmid expressing GFP, and enhancer activity was observed after 4 days (Figure 7D). Indeed, human CRE6 was able to drive reporter gene expression in human cortical organoids (Figure 7E). Comparing hCRE6 and mCRE6, higher levels of mScarlet were observed for hCRE6 compared to the orthologous mouse sequence (Figure 7E). Human CRE9 was also able to drive mScarlet expression at similar levels as hCRE6, yet, in this case the orthologous mouse sequence showed similar or even higher activity (Figure 7F). The enhancer activity assay therefore confirms that at least two of the human putative CREs identified based on their open chromatin configuration have the potential to enhance gene expression from a reporter plasmid. For the endogenous locus, that is embedded in chromatin, the epigenetic state of the locus may further contribute to tuning the activity of the CRE.

Thus, based on the open chromatin status, which correlates with gene expression in the hNcx, and closed chromatin correlating with repression in the mNcx, respectively, these 11 genomic regions may represent putative enhancers contributing to differential *EREG* expression in the developing neocortex of different mammalian species (Figure S7).

## DISCUSSION

Here, we show that the growth factor EPIREGULIN provides a pro-proliferative signal in the human SVZ niche that is received by BPs and promotes their amplification. EPIREGULIN is expressed in apical and basal radial glia of the human neocortex, where it is required for basal progenitor proliferation. EPIREGULIN is also expressed in macaque and gorilla, both gyrencephalic primates, but is not detectable in the mouse, a species with a small lissencephalic neocortex. In the mNcx, addition of EPIREGULIN promotes BP amplification, of both bIPs and bRG. This may have important evolutionary relevance as BPs have high self-renewing capacities, are highly abundant in species with a large gyrencephalic neocortex and are associated with increased neocortex size, neuron number and cortical folding (Dehay et al., 2015; Florio and Huttner, 2014; Llinares-Benadero and Borrell, 2019; Lui et al., 2011; Zhou et al., 2023). Human mutations affecting BP generation and amplification often result in microcephaly (Baala et al., 2007; Johnson et al., 2018). Further addition of EPIREGULIN to the hNcx did not increase BP proliferation to even higher levels, suggesting that the system may have been saturated, either at the level of receptors (see below) or that maximum BP proliferation had already been reached. As addition of EPIREGULIN to the gorilla Ncx resulted in increased BP proliferation, this suggests that the impact of EPIREGULIN on BP amplification depends on the prevailing EPIREGULIN concentration in a given species and/or on the species-specific context, as reported for other genes (Van Heurck et al., 2022), and may provide a mechanism to tune progenitor amplification across species.

Based on transcriptomic studies, it was proposed that bRG maintain the OSVZ as a proliferative stem cell niche by locally producing growth factors and ECM to activate self-renewal pathways (Fietz et al., 2010; Kalebic and Huttner, 2020; Pollen et al., 2015). In particular, the germinal zones, including the mVZ, hVZ, hISVZ and hOSVZ, but not the mSVZ, preferentially express genes related to ECM formation (Fietz et al., 2010; Florio et al., 2015; Pollen et al., 2015). Indeed, targeted activation of the ECM receptor integrin αvβ3 on BPs in the mNcx promoted BP expansion (Stenzel et al., 2014). Moreover, conditional expression of Sox9, a transcription factor that induces expression of ECM components, resulted in cell-non-autonomous stimulation of BP proliferation linked to increased neuron production (Guven et al., 2020). In addition to proliferation, specific ECM components with higher expression in human than mouse NPCs were shown to cause folding of the hNcx (Long et al., 2018). Here, we provide functional evidence that a growth factor expressed in primate bRG promotes BP amplification in the SVZ of the mNcx and contributes to BP proliferation in the hNcx. In gyrencephalic species, BPs were shown to have a higher number of processes enabling them to receive extrinsic pro-proliferative signals linked to an increased BP proliferative capacity (Kalebic et al., 2019). Our data suggests that EPIREGULIN provides such a pro-proliferative signal in the human SVZ niche and stimulates BPs amplification.

Moreover, transcriptional signatures of bRG have been reported to be enriched in cells from human primary glioblastoma, suggesting that developmental programs are reactivated in tumor cells (Bhaduri et al., 2020; Pollen et al., 2015). Indeed, *EREG* expression is higher in high-compared to low-grade glioma and loss of *EREG* correlates with increased survival. This is in line with the induction of NPC proliferation by EPIREGULIN described here, even though EPIREGULIN may also contribute to other aspects of tumor progression, such as angiogenesis or inflammation (Riese and Cullum, 2014).

Mechanistically, the addition of EPIREGULIN did not appear to alter transcriptional programs but rather resulted in altered cell type proportions, as has also been observed for other factors that regulate brain size, such as abnormal spindle-like microcephaly-associated (*ASPM*), the most common recessive gene associated with human primary microcephaly (Johnson et al., 2018). Both the mNSC proliferation assay in the presence/absence of EGF and the inhibition of EGFR suggest that EPIREGULIN mediates NPC proliferation, resulting in altered cell type proportions, via EGFR mediated signaling. Among the main signaling cascades activated by EGFR ligand binding are the Ras-Raf-MEK-ERK, STAT and PI3K-AKT-mTOR pathways, all of which are important for the regulation of NPC proliferation, differentiation and progenitor pool maintenance (Kalebic and Huttner, 2020; Pollen et al., 2015; Romano and Bucci, 2020). In contrast to EGF, which represents a high affinity ligand reported to bind cell-surface EGFR with an apparent Kd of 0.1–1 nM, EPIREGULIN represents a low affinity ligand with 10- to 100-folder weaker binding. Interestingly, EGF was reported to promote transient EGFR activation resulting in transient ERK activation, whereas the low affinity ligand EPIREGULIN was counterintuitively shown to promote both sustained EGFR activation and sustained ERK activation (Freed et al., 2017). The distinct biological responses were independent of any effects on other ErbB family receptors, which is in line with our finding that both specific EGFR inhibition and pan-ErbB inhibition resulted in a similar loss of the EPIREGULIN-mediated increase in basal mitosis. Thus, ErbB4 does not appear to play a major player in the EPIREGULIN response in the mNcx, despite its high expression in BPs.

The stimulation of BP amplification by EPIREGULIN was dose-dependent, reaching a maximum at 50 ng/mL in the mNcx. At 100 ng/mL, EPIREGULIN did not induce any increase in proliferation in the mNcx. Similarly, human BPs, in contrast to gorilla BPs, were not receptive to further stimulation of proliferation by EPIREGULIN. This suggests that levels of individual growth factors and their combinations present in the tissue dictate cellular behavior. Moreover, different desensitization strategies have been reported for ErbB receptors and their distinct ligands, which may further attenuate the response to high levels of those ligands (Yamamoto et al., 2014).

Research has only begun to unravel the genomic basis for neocortex evolution. Recently, a handful of human-, hominid- and primate-specific genes have emerged as potential adaptive evolutionary innovations implicated in neocortex evolution (Florio et al., 2017; Libe-Philippot and Vanderhaeghen, 2021; Pollen et al., 2023). However, species-specific differences are likely to extend beyond novel genes and there is compelling evidence emerging that non-coding regions of the genome contribute to evolutionarily relevant inter-species differences in gene expression (Doan et al., 2018; Silver, 2016). Among the few examples of regulatory changes that were functionally investigated is *HARE5*, a human-accelerated regulatory enhancer that drives earlier and more robust brain expression of *FZD8*, a WNT receptor, resulting in accelerated cell cycle and enlarged brains (Boyd et al., 2015). Moreover, deletion of a 15 base pair region in a regulatory element of *GPR56* selectively influences NPC proliferation and neocortex folding, potentially implicated in neocortex evolution (Bae et al., 2014). Here, we have identified CRE regions adjacent to human *EREG* that are marked by open chromatin suggesting that they may represent putative active enhancers. Indeed, the two hCRE regions that we tested in enhancer activity assays were able to promote reporter gene expression. The homologous mouse regions could be identified for 10 of the 11 regions, yet show higher sequence divergence compared to other primates and lack signs of active chromatin. Of the two mCREs that we tested in enhancer activity assays, one showed reduced activity compared to human, whereas the other sequence showed comparable activity. Given that the endogenous CRE regions are embedded in chromatin, local epigenetic modifications and chromatin compaction may further contribute to regulating the accessibility of factors and, thus, the activity of the regions. Overall, these potential CREs represent interesting candidate enhancers that may contribute to inter-species differences in *EREG* expression.

Taken together, we propose that EPIREGULIN represents a pro-proliferative signal for BP cells in primates to sustain a proliferative niche in the subventricular zone of the developing neocortex, which may have important evolutionary significance as a tunable mechanism to control neocortex size across mammalian species.

## ACKNOWLEDGEMENTS

We are grateful to the facilities of the CRTD and Dresden Concept partners for the outstanding support provided, notably K. Neumann and her team of the Stem Cell Engineering Facility, H. Hartmann and her team of the Light Microscopy Facility, A. Gompf and her team of the Flow Cytometry Facility, the teams for animal husbandry and histology, the Dresden Concept Genome Center lab team for technical support and the MPI-CBG Genome Engineering Facility. We thank all members of the Albert laboratory for help and discussions. We thank I. Mestres and A. Pinson for advice on the IUE, F. Noack for advice on the enhancer activity analysis and the Bonev lab for kindly providing the construct for the enhancer activity assay. This work was supported by the Center for Regenerative Therapies TU Dresden, the DFG (Emmy Noether, AL 2231/1-1), the Schram foundation, and ERA-NET Neuron (MEPIcephaly). Specifically, the topic on which this report is based was funded by the Federal Ministry of Education and Research (BMBF) under the funding code 01EW2208. The content of this publication is the responsibility of the authors. Moreover, this project was funded by the Federal Ministry of Education and Research (BMBF) and the Free State of Saxony as part of the Excellence Strategy of the federal and state governments, in the frame work of transCampus.

## AUTHOR CONTRIBUTIONS

Conceptualization, P.C. and M.A.; Investigation, P.C., N.D., A.K., G.S., J.E., T.M.S. and M.A.; Resources, A.K., N.D., J.E., R.P.D, C.B., J.E.M.K., M.S., U.M., K.L. and P.W.; Formal Analysis, P.C., N.D., A.P.; Visualization, P.C., N.D.; Writing – Original Draft, M.A.; Writing – Review & Editing, P.C., N.D., A.K., G.S., J.E., T.M.S., K.R.L. and M.A.; Funding Acquisition, K.R.L. and M.A.; Supervision, M.A.

## METHODS

### Mice

All experimental procedures were conducted in agreement with the German Animal Welfare Legislation after approval by the Landesdirektion Sachsen (licenses DD24.1-5131/476/8; 25-5131/496/19). Animals used for this study were kept in standardized hygienic conditions with free access to food and water and on a 12-hour/12-hour light/dark cycle. All mice were wildtype mice from the inbred C57BL/6J strain, except for the isolation of neural cell populations for RNA-seq, where the transgenic *Tubb3*::GFP line (Attardo et al., 2008) was used to identify neurons based on GFP expression. Embryonic day (E) 0.5 was set as noon on the day on which the vaginal plug was observed. All experiments were performed in the dorsolateral telencephalon of mouse embryos, at a medial position along the rostro-caudal axis. The developmental time point E14.5 of experimental procedures corresponds to a mid-neurogenic stage, when the production of upper-layer neurons has started. The sex of embryos was not determined, as it is not likely to be of relevance for the results obtained in the present study. Mouse NSCs were isolated from E11.5/12.5 embryos as previously described (Schmitz et al., 2011; Schütze et al., 2023).

### Human fetal brain tissue

Human fetal brain tissue was obtained from the Department of Gynecology and Obstetrics, University Clinic Carl Gustav Carus of the Technische Universität Dresden, following elective pregnancy termination and informed written maternal consents, and with approval of the local University Hospital Ethical Review Committee (ethical approval number EK 355092018). The age of fetuses ranged from gestation week (GW) 10 to 13 as assessed by ultrasound measurements of crown-rump length and other standard criteria of developmental stage determination. These developmental time points correspond to an early/mid-neurogenic stage, when the OSVZ expands and the production of upper-layer neurons starts. Due to protection of data privacy, the sex of the human fetuses, from which the hNcx tissue was obtained, cannot be reported. The sex of the human fetuses is not likely to be of relevance for the results obtained in the present study. No health disorders were reported for any of the fetal hNcx tissue samples used in this study. Fetal human brain tissue was dissected in Tyrode’s solution and used immediately (within 1 h) for further culture or fixation.

### Human and primate induced pluripotent stem cell lines

All experiments involving hiPSCs were performed in accordance with the ethical standards of the institutional and/or national research committee, as well as with the 1964 Helsinki Declaration and its later amendments, and approved by the University Hospital Ethical Review Committee (ethical approval number SR-EK-456092021). Human cortical organoids were generated using the previously generated human induced pluripotent stem cell (iPSC) line CRTDi004-A (https://hpscreg.eu/cell-line/CRTDi004-A) (Völkner et al., 2022), derived from a healthy donor. The human iPSC line was maintained on Matrigel-coated (Corning, 354277) culture dishes in mTeSR1 and passaged using TrypLE Express enzyme (Schütze et al., 2023).

Gorilla organoids were created using the previously generated gorilla iPSC line GC1 (Wunderlich et al., 2014). This line was maintained on Matrigel-coated culture dishes in StemFlex™ Medium and passaged using ReLeSR.

### *Ex vivo* experiments on mNcx

Mouse organotypic slice cultures were performed from E14.5 brains, as previously described (Wong et al., 2014). The meninges were surgically removed from brains, brains were embedded in 3% low melting point agarose (Invitrogen, 16520100) and cut on a vibratome into 250 μm-thick slices. Slices were embedded into a collagen I-based hydrogel and placed in 35 mm Petri dishes with a 14 mm microwell. Slice culture medium, either without (control) or with 10 ng/mL, 50 ng/mL or 100 ng/mL human recombinant EPIREGULIN, dissolved in PBS, were added to the slices. Slice culture medium (SCM) contained Neurobasal medium supplemented with 10% rat serum, 20 μM glutamine, 1X penicillin/streptomycin, N-2 and B-27 supplement and 0. 1 M HEPES/NaOH, pH 7.2. Slices were cultured for 24 hours in a humidified incubation chamber supplied with 40% O_2_, 5% CO_2_ and 55% N_2_ at 37°C.

Mouse brain hemispheres were cultured under rotation conditions (HERO), as previously described (Schenk et al., 2009). The meninges were surgically removed from E14.5 brains, hemispheres dissected and placed into rotating flasks with 1.5 mL of SCM. SCM was either without (control) or with 50 ng/mL human recombinant EPIREGULIN. HERO culture was performed for 24 hours in a Whole Embryo Culture System (Nakayama, 10-0310) in a humidified atmosphere of 40% O_2_, 5% CO_2_ and 55% N_2_ at 37°C.

### Mouse NSC culture

Mouse NSCs were plated on 6-well plates coated with poly-D-lysine and laminin at a density of 105 cells/mm^2^ and grown in NSC medium which contains a 1:1 mixture of DMEM/F12 and Neurobasal medium supplemented with 10 ng/mL epidermal growth factor (EGF), 20 ng/mL fibroblast growth factor (FGF), N-2 supplement, B-27 supplement, Penicillin/Streptomycin, Sodium-Pyruvate, GlutaMAX, MEM-NEAA, Heparin, HEPES, 2-mercaptoethanol and BSA (Schmitz et al., 2011; Schütze et al., 2023). NSCs are passaged using trypsin-EDTA and soybean trypsin inhibitor and kept in standard conditions at 37°C and 5% CO_2_.

### *Ex vivo* experiments on hNcx

Fetal hNcx tissue was cultured under free-floating tissue culture (FFTC) conditions, as previously described (Long et al., 2018). Fetal hNcx tissue was placed into 1.5 mL of SCM, either without (control) or with 50 ng/mL human recombinant EPIREGULIN, and cultured for 24 hours under rotation conditions in a humidified atmosphere of 40% O_2_, 5% CO_2_ and 55% N_2_ at 37°C.

### Generation of human and gorilla organoids

Human and gorilla cortical organoids were generated according to the sliced neocortical organoid (SNO) method, as previously described in detail (Qian et al., 2020; Schütze et al., 2023). Briefly, hiPSC colonies of about 1.5 mm in size were detached with collagenase and transferred to ultra-low attachment 6-well plates (Corning, 3471), containing forebrain medium 1, to form 3D aggregates. On day 7, embryoid bodies (EBs) were embedded in Matrigel (Corning, 354230) with about 20 EBs per Matrigel cookie. From day 7 to 14, EBs were cultured in forebrain maturation medium 2. At day 14, organoids were mechanically released from the Matrigel cookie and further cultured in 6-well ultra-low attachment plates in forebrain medium 3 on an orbital shaker at 100 rpm. From day 35, 1% v/v Matrigel was added to forebrain medium 3. At 6 weeks, organoids were embedded in 3% low melting point agarose and cut on a vibratome into 500 μm-thick slices, supporting enhanced supply of oxygen and nutrients throughout the organoids. Slicing of organoids was repeated at 10 weeks. From day 72, organoids were cultured in forebrain medium 4. For treatments, 50 ng/mL of EPIREGULIN were added directly after slicing of organoids, either at 6 weeks or 10 weeks, to avoid issues with penetration. EPIREGULIN was added every other day for a period of 4 or 7 days (6-week organoids) or 10 days (10-week organoids).

Additionally, gorilla cerebral organoids were generated according to the cerebral organoid method, using the STEMdiff Cerebral Organoid Kit, based on (Lancaster and Knoblich, 2014; Lancaster et al., 2013).

### Guide RNA design and validation

For CRISPR/Cas9-mediated targeting of the human *EREG* gene, three guide RNAs (gRNAs; see Table S1) were designed using Geneious. The previously published gRNA for *LacZ* (Kalebic et al., 2016; Platt et al., 2014) was used as control. Following Integrated DNA Technologies (IDT) Alt-R CRISPR-Cas9 System, gRNAs were purchased as custom crRNAs and assembled into functional gRNA duplexes with the tracrRNA according to the manufacturer’s instructions. Ribonucleoprotein (RNP) complexes were assembled using 24 μM gRNA and 8 μM recombinant Cas9 protein from *Streptococcus pyogenes* (1:3 ratio Cas9:gRNA). The gRNAs g*EREG* KO1 and g*EREG* KO2 were assembled as separate RNPs, and then used as 1:1 mix.

Targeting of the gRNAs was validated using two different systems. Firstly, *in vitro* cleavage of target DNA was carried out. In brief, to create the template, PCR was performed with 2x NEBNext High-Fidelity PCR Master Mix (New England Biolabs, M0541S) and 10 μM primers (Table S2) on genomic DNA isolated from CRTDi004-A iPSC under the following cycling conditions: initial denaturation at 98°C for 30 sec, 40 cycles [98°C 10 sec, 58/60°C 30 sec, 72°C 1 min] and final elongation at 72°C for 5 min. PCR products were analyzed via gel electrophoresis on a 1.5% agarose gel, cut from the gel and extracted using the QIAquick gel extraction kit. Templates were combined with the generated RNP complexes, incubated for 60 mins at 37°C, the digestion stopped with 2 mg/mL Proteinase K (56°C, 10 min) and the final cleavage products analyzed on a 2% agarose gel.

Secondly, gRNAs were validated in CRTDi004-A iPSCs. For this, 160,000 cells per condition were nucleofected with the RNPs and 0.1 μg/μl of a pCAG-GFP plasmid using the 4D-Nucleofector by Lonza (CB150 pulse, P3 Primary Cell 4D-Nucleofector X Kit S; Lonza, V4XP-3032). Nucleofected cells were plated on Matrigel-coated 6-well plates and cultured in mTeSR1 with 1x Pen/Strep. After three days in culture, cells were detached using TrypLE Express Enzyme, resuspended in 300 μl of 2% FBS in PBS with 1000 ng/μl DAPI and processed by FACS on a Melody Cell Sorter (BD Biosciences) using the BD FACSChorus software. Roughly 10,000–60,000 GFP-positive cells per condition were collected in 2% FBS in PBS and DNA extraction was performed. PCR templates were generated as described above using Seq-Primers (Table S2), gel-purified and sent for Sanger sequencing (Microsynth) using the forward primers (Table S2). The sequencing results were analyzed using Geneious Prime (pairwise, global alignment with free end gaps, 93% similarity cost matrix).

### CRISPR/Cas9-mediated gene knockout in human cortical organoids

Electroporation of the RNP complexes for CRISPR/Cas9-mediated targeting of *EREG* was carried out 2 days after the first slicing of human cortical organoids at week 6 of organoid culture. Organoids were transferred to a 6 cm ultra-low-attachment dish (Eppendorf, 30701011) containing Tyrode’s solution (Sigma, T2145). Using a glass microcapillary (Sutter Instrument, BF120-69-10), 0.2 to 0.5 μl of the RNP/plasmid (pCAG-GFP) mix at a final concentration of 24 μM RNP and 0.1 μg/μl plasmid diluted in 0.1% Fast Green Solution (in water) were injected into areas depicting ventricular morphology. Injections were carried out using a microinjector (World Precision Instruments, SYS-PV820) on continuous setting. Up to five ventricles were injected per organoid. A total of 7-8 organoids were processed per replicate. After injection, organoids were transferred into an electroporation chamber containing Tyrode’s solution and electroporated with 5 pulses applied at 38 V for 50 ms each at intervals of 1 s (Harvard Bioscience, BTX ECM 830). Subsequently, the electroporated organoids were moved back into culture medium and incubated for 7 days before fixation with 4% PFA.

### Immunohistochemistry

Tissue was fixed in 4% PFA in 120 mM phosphate buffer pH 7.4 for 24 hours at 4°C, transferred to 30% sucrose for 24 hours, embedded in O.C.T. compound (Sakura Finetek, 4583) with 15% sucrose and cut into 12 μm (human tissue), 20 μm (mouse tissue) or 25-30 μm (organoids) cryosections. For analysis of cell morphology (Figure 3I–M), fixed tissue was cut into 70 μm vibratome sections.

Immunofluorescence was performed as previously described (Florio et al., 2015). Antigen retrieval for 1 hour with 10 mM citrate buffer pH 6.0 at 70°C in a water bath was followed by a wash with PBS, quenching for 30 min in 0.1 M glycine in PBS and blocking for 30 min in blocking solution (10% horse serum and 0.1% triton in PBS; or glycine 0.2%, 300 mM NaCl and 0.3% triton in PBS for phospho-vimentin staining) at room temperature. Primary antibodies were incubated in blocking buffer over night at 4°C. Subsequently, sections were washed 3 times in PBS, incubated with secondary antibodies (1:1,000) and DAPI (1:1,000) in blocking solution for 1-2 hours at room temperature, and washed again 3 times in PBS before mounting on microscopy slides with Mowiol.

Most images were acquired using either with a Zeiss LSM 980 confocal microscope with a 20x objective using 0.83-μm or 0.50-μm thick optical sections or with a 40x objective with 0.34-μm optical sections. Few images were acquired either with a Keyence BZ-X800E fluorescence microscope with a 10x objective or with a Zeiss ApoTome2 fluorescence microscope with a 20x objective and 1.5-μm thick optical sections. Images are shown as maximum intensity projection of 11-15 optical sections. When images were taken as tile scans, the stitching of tiles was performed using the ZEN software.

### Gene expression analysis by RT-qPCR

Gene expression analysis was performed as previously described (Albert et al., 2017). For RT-qPCR, RNA was isolated from 10,000 cells using the RNeasy Mini kit, cDNA was synthesized using random hexamers and Superscript III Reverse Transcriptase (Invitrogen) and qPCR performed with LightCycler 480 SYBR Green I Master (Roche) on a Light Cycler 480 (Roche). For each sample, three technical replicates were run. Gene expression data was normalized based on the housekeeping gene *Gapdh*. Primers are listed in Table S2.

### Isolation of neural cell populations for RNA-seq

Isolation of aRG, bIPs and neurons has been previously described (Florio et al., 2015; Molyneaux et al., 2015; Schütze et al., 2023). After HERO culture of E14.5 cerebral hemispheres from *Tubb3*::GFP mice (Attardo et al., 2008) with or without 50 ng/mL EPIREGULIN for 24 hours, the dorsolateral neocortex was dissected and a single cell suspension was produced from 2-3 hemispheres per condition using the MACS Neural Tissue Dissociation kit containing papain (Miltenyi Biotec, 130-092-628). Cells were fixed in 1% formaldehyde for exactly 10 min on a rotating wheel (10 rpm) at room temperature, the fixative was quenched by the addition of 0.2 M glycine for 5 min, cells were washed in 1% BSA with centrifugation for 5 min at 500 × g, permeabilized for 10 min at 4°C with 0.1% saponin (Sigma Aldrich, 47036) and 0.2% BSA (Sigma Aldrich, A2153) in Tyrode’s solution and subsequently spun down for 3 min at 500 × g. At least 3 million cells were stained with antibodies against Sox2 (Mouse anti-Sox2-PE, 1:40) and Tbr2 (Rabbit anti-TBR2/Eomes, 1:250), washed twice in permeabilization buffer and once in 0.5% BSA. This was followed by staining with an anti-rabbit secondary Alexa405 (1:1000). The cells were resuspended to 1–3 million cells/100 μL in Tyrode’s solution, passed through a 35 μm filter and immediately processed by FACS on a 5-laser FACS Aria II sorter using the FACS Diva software. For each cell type and condition, 250,000 cells were collected in 1.5 mL DNA LoBind tubes containing lysis buffer. RNA was extracted using the Quick-RNA FFPE MiniPrep kit. Biological replicates (n=3) were collected from three pregnant female mice that were processed and sorted independently on different days.

### RNA-seq library preparation

Transcriptome libraries were prepared using the SmartSeq 2 protocol (Picelli et al., 2013). Isolated total RNA from an equivalent of 25,000 cells was denatured for 3 minutes at 72°C in 4 μL hypotonic buffer (0.2% Triton-X 100) in the presence of 2.5 mM dNTP, 100 nM dT-primer and 4 U RNase Inhibitor (Promega, N2611). Reverse transcription was performed at 42°C for 90 min after filling up to 10 μl with RT buffer mix for a final concentration of 1x superscript II buffer (Invitrogen), 1 M betaine, 5 mM DTT, 6 mM MgCl2, 1 μM TSO-primer (Table S2), 9 U RNase inhibitor and 90 U Superscript II. The reverse transcriptase was inactivated at 70°C for 15 min. For subsequent PCR amplification of the cDNA the optimal PCR cycle number was determined with an aliquot of 1 μL unpurified cDNA in a 10 μL qPCR containing 1X Kapa HiFi Hotstart Readymix (Roche), 1x SybrGreen and 0.2 μM UP primer (Table S2). The residual 9 μL cDNA were subsequently amplified using Kapa HiFi HotStart Readymix (Roche) at a 1x concentration together with 250 nM UP-primer (Table S2) under the following cycling conditions: initial denaturation at 98°C for 3 min, 12 cycles [98°C 20 sec, 67°C 15 sec, 72°C 6 min] and final elongation at 72°C for 5 min. Amplified cDNA was purified using 1x volume of Sera-Mag SpeedBeads (GE Healthcare) resuspended in a buffer consisting of 10 mM Tris, 20 mM EDTA, 18.5% (w/v) PEG 8000 and 2 M sodium chloride solution. The cDNA quality and concentration were determined using a Fragment Analyzer (Agilent).

For library preparation, 2 μl amplified cDNA was tagmented in 1x Tagmentation Buffer using 0.8 μl bead-linked transposome (Illumina DNA Prep, (M) Tagmentation, Illumina) at 55°C for 15 min in a total volume of 4 μL. The reaction was stopped by adding 1 μl of 0.1% SDS (37°C, 15 min). Magnetic beads were bound to a magnet, the supernatant was removed, beads were resuspended in 14 μl indexing PCR Mix containing 1x KAPA Hifi HotStart Ready Mix (Roche) and 700 nM unique dual indexing primers (i5 and i7), and subjected to a PCR (72°C 3 min, 98°C 30 sec, 12 cycles [98°C 10 sec, 63°C 20 sec, 72°C 1 min], 72°C 5 min). Libraries were purified with 0.9x volume Sera-Mag SpeedBeads, followed by a double size selection with 0.6x and 0.9x volume of beads. Sequencing was performed after quantification using a Fragment Analyzer on an Illumina Novaseq 6000 with an average sequencing depth of 60 million fragments.

### Histone methylation analysis by ChIP-qPCR

ChIP was performed as previously described (Albert et al., 2017). For ChIP-qPCR, 100,000 cells were sorted into 1.5 mL tubes, filled up to 500 μL with PBS, and transferred into DNA LoBind tubes. Cells were fixed in 1% formaldehyde while shaking at 700 rpm for 10 min at room temperature. The reaction was stopped by addition of 80 μL 1.25 M glycine and incubated at 350 rpm for 5 min at RT. To facilitate pelleting of cells, 60 μL fetal bovine serum was added (Adli and Bernstein, 2011), and cells were centrifuged at 1,600 × g for 5 min at 4 °C. Cells were washed twice with 1 mL PBS containing 10% FBS and then resuspended in 80 μl cold lysis buffer containing 1% SDS, 50 mM Tris-HCl (pH 8.1), 100 mM NaCl, 5 mM EDTA, and protease inhibitors and incubated on ice for 5 min with intermittent brief vortexing. The samples were sonicated on a Covaris S2 in microTUBES with AFA Snap cap for 3 min with 2% duty cycle, intensity of 3,200 cycles per burst using frequency sweeping as power mode and continuous degassing at 4°C. ChIP was performed using the LowCell ChIP kit according to manufacturer’s instructions. For each ChIP, 1 μL of anti-H3K27me3 (Cell Signaling, 9733) was used. For the qPCR the LightCycler 480 SYBR Green I Master and the Light Cycler 480 from Roche were used. Primers are listed in Table S2.

### Constructs for epigenome editing

The constructs for epigenome editing were based on the plasmid pSpCas9n(BB)-2A-GFP (Addgene #48140) (Ran et al., 2013) for expression of Cas9 and GFP, linked by a T2A site, from a CAGS promoter. Modifications included conversion of the Cas9 nickase into a non-cutting dCas9, introduction of a BsaI-flanked ccdB cassette between the U6 promoter and guide RNA scaffold, and introduction of two BbsI sites between the C-terminus of dCas9 and the NLS sequence. The coding sequence of the catalytic domain of human KDM6B (1025–1680 aa) was obtained from the MSCV_JMJD3 plasmid (Addgene #21212) (Sen et al., 2008). BsaI sites in the coding sequence were inactivated by introduction of silent mutations through PCR amplification of KDM6B fragments and reassembly downstream of dCas9. The resulting plasmid backbone pCAGS_dCas9-KDM6B-L_GFP_ccdB featured sequences for a dCas9-JMJC_6B fusion protein and GFP expressed from a CAGS promoter as well as an U6 promoter followed by a BsaI-ccdB cassette for efficient gRNA cloning.

Four gRNAs targeting sequences upstream and downstream of the mouse *Ereg* transcription start site were selected using Geneious Prime software (see Table S1 for gRNA sequences). Two pairs consisting of an upstream and a downstream gRNA expressed from the human U6 promotor and fused to a gRNA scaffold were cloned into the dCas9-JMJC_6B-T2A-EGFP plasmid. Sequences encoding target specific gRNAs were cloned by golden Gate cloning using BsaI. The ccdB cassette was excised and replaced by a cassette featuring the spacer and scaffold for the first gRNA followed by a second U6 promoter and a second spacer. CcdB removal allowed counter selection. The previously published gRNA targeting *LacZ* was used as control (Albert et al., 2017; Kalebic et al., 2016).

### Epigenome editing in mNSC

For epigenome editing, mouse NSC were transfected with the dCas9-JMJC_6B-T2A-EGFP plasmid encoding a pair of gRNAs targeting the *Ereg* locus. Two to three million NSCs were resuspended in 100 μL P3 primary solution (Lonza) containing 10 μg of the plasmid DNA and transfected using the program DS-112 of the 4D-nucleofector (Lonza). NSCs were harvested 48 hours post-nucleofection, resuspended in Tyrode’s solution containing 0.5% BSA and GFP-positive cells collected by FACS. Cells were then further processed for RT-qPCR and ChIP-qPCR.

### Enhancer activity assay

The constructs for examining enhancer activity of human candidate regulatory regions and their mouse orthologues were generated as previously described (Noack et al., 2022) and a detailed protocol can be found at https://www.protocols.io/view/mpra-plasmid-pool-preparation-bxchpit6/. Briefly, 312-bp single-stranded oligonucleotides containing candidate regulatory elements or scrambled control sequences and overhangs for Gibson assembly were synthesized (Twist Bioscience) (all sequences are listed in Table S3). These inserts were introduced into the pMPRA1 Addgene plasmid backbone (no. 49349) (Melnikov et al., 2014) via Gibson assembly and transformed into ElectroMAX Stbl4 Competent Cells (ThermoFisher, 11635018) using 1.8 kV, 25 μF and 200 Ω. The purified plasmids were digested with KpnI/EcoRI and ligated with an insert containing a minimal promoter and mScarlet-I (kindly provided by the Bonev Lab). The purified ligation product was heat-transformed in *Escherichia coli*. The final plasmids were purified using the GeneJET Plasmid-Midiprep-Kit (Thermo Scientific, K0481). All primers used are listed in Table S2.

The plasmids were co-electroporated with a CAG-GFP plasmid (ratio of pMPRA1:CAG-GFP is 3:1) into human cortical organoids at week 6 as described above. Three days post electroporation the organoids were fixed and immunohistochemistry was performed for GFP and for mScarlet using the anti-RFP antibody, as described above.

### RNA-seq data analysis

Basic quality control of the sequence data was done with FastQC (v0.11.6). Adapters (nextera:CTGTCTCTTATA) and polyA/T tails were trimmed with cutadapt (v2.6) (Martin, 2011) and pairs where one or both reads were shorter than 35 bp were removed. Trimmed reads were aligned to the mouse reference genome GRCm39 using the aligner gsnap (v2020-12-16) (Wu and Nacu, 2010) with Ensembl 104 mouse splice sites as support. Uniquely mapped reads were compared based on their overlap to Ensembl 104 human gene annotations using featureCounts (v2.0.1) (Liao et al., 2014) to create a table of fragments per mouse gene and sample. Normalization of raw fragments based on library size and testing for differential expression between the different cell types/treatments was performed with the R package DESeq2 (v1.30.1) (Love et al., 2014). Principal component analysis based upon the top 500 genes with the highest variance were computed to explore correlation between biological replicates and different libraries. To identify differentially expressed genes, counts were fitted to the negative binomial distribution and genes were tested between conditions using the Wald test of DESeq2. Resulting p-values were corrected for multiple testing with the Independent Hypothesis Weighting package (IHW 1.18.0) (Ignatiadis and Huber, 2017; Ignatiadis et al., 2016). Genes with a maximum of 10% false discovery rate (padj ≤ 0.1) were considered as significantly differentially expressed. TPM values for quantifying gene expression levels within samples were calculated with kallisto (v0.46.1) (Bray et al., 2016) based on an index of Ensembl 104 mouse transcript sequences and TPMs were summarized at gene level. RNA-seq data has been deposited with the Gene Expression Omnibus under accession code GSE228007 (reviewer access token atixosskplqzlgb).

### Cell counting

All samples were blinded before the acquisition of the data. All cell counts were performed in standardized microscopic fields using Fiji, processed using Excel (Microsoft), and results were plotted using Prism (GraphPad Software). Image quantifications were performed either manually or using the Fiji plugin Stardist (Schmidt et al., 2018). Shortly, images were analyzed as maximum intensity projections. Images were cropped to either 400 μm (PH3), 200 μm (Sox2, Tbr2, Ki67 and PCNA) or 100 μm (IUE) wide pictures, followed by crop of the different cortical layers (VZ, SVZ/IZ, CP). Channels were split and the Stardist 2D plugin was performed using the versatile (fluorescent nuclei) model. If necessary, segmentation was corrected manually. Abventricular PH3 was defined as more than three nuclei away from the ventricular surface using DAPI as reference. Quantification of electroporated samples was performed by segmenting the GFP cells either with Stardist or manually, and then counting double-positive cells using the cell counter tool in Fiji. The proliferation assay in mNSC was quantified using the BZ-X800 analyzer software.

### Statistical analysis

Sample sizes are reported in each figure legend. All statistical analysis was performed using Prism (GraphPad Software). Normal distribution of datasets was tested by Shapiro-Wilk or Kolmogorov-Smirnov test. The tests used included Student’s *t*-test and one-way ANOVA with Dunnett post hoc test, as indicated in the figure legend for each quantification. Significant changes are indicated by stars for each graph and described in the figure legends.

## RESOURCE TABLE

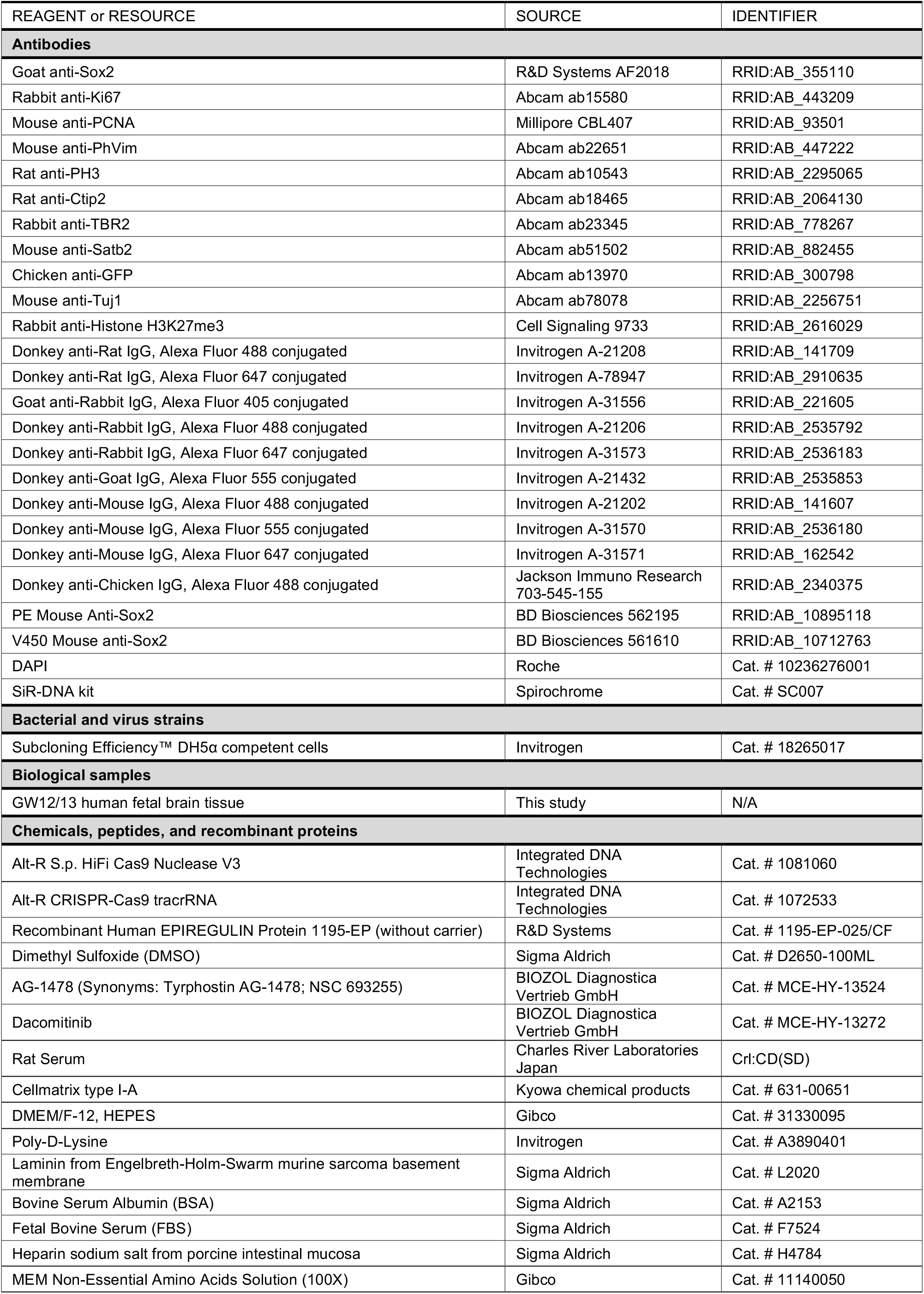
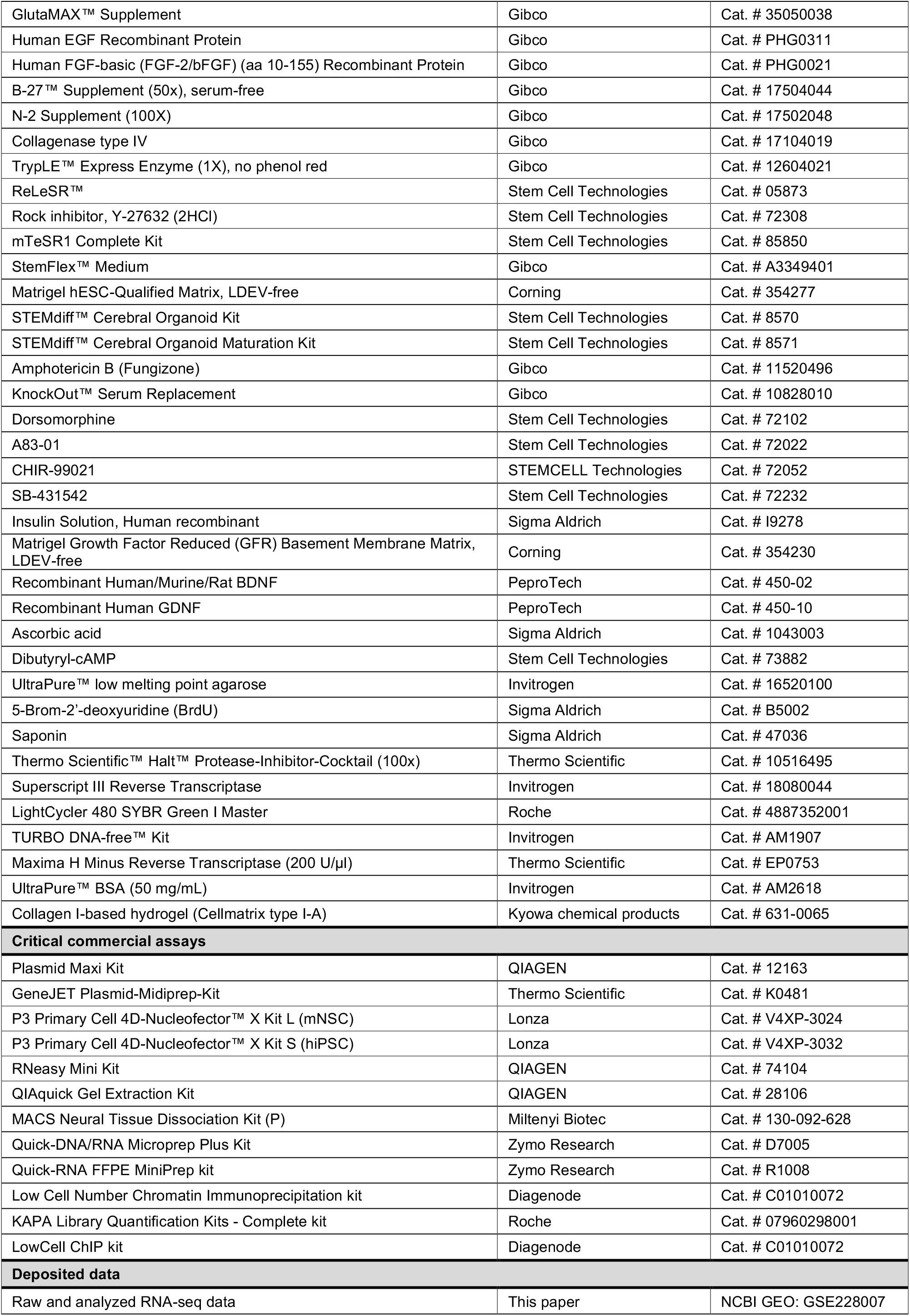
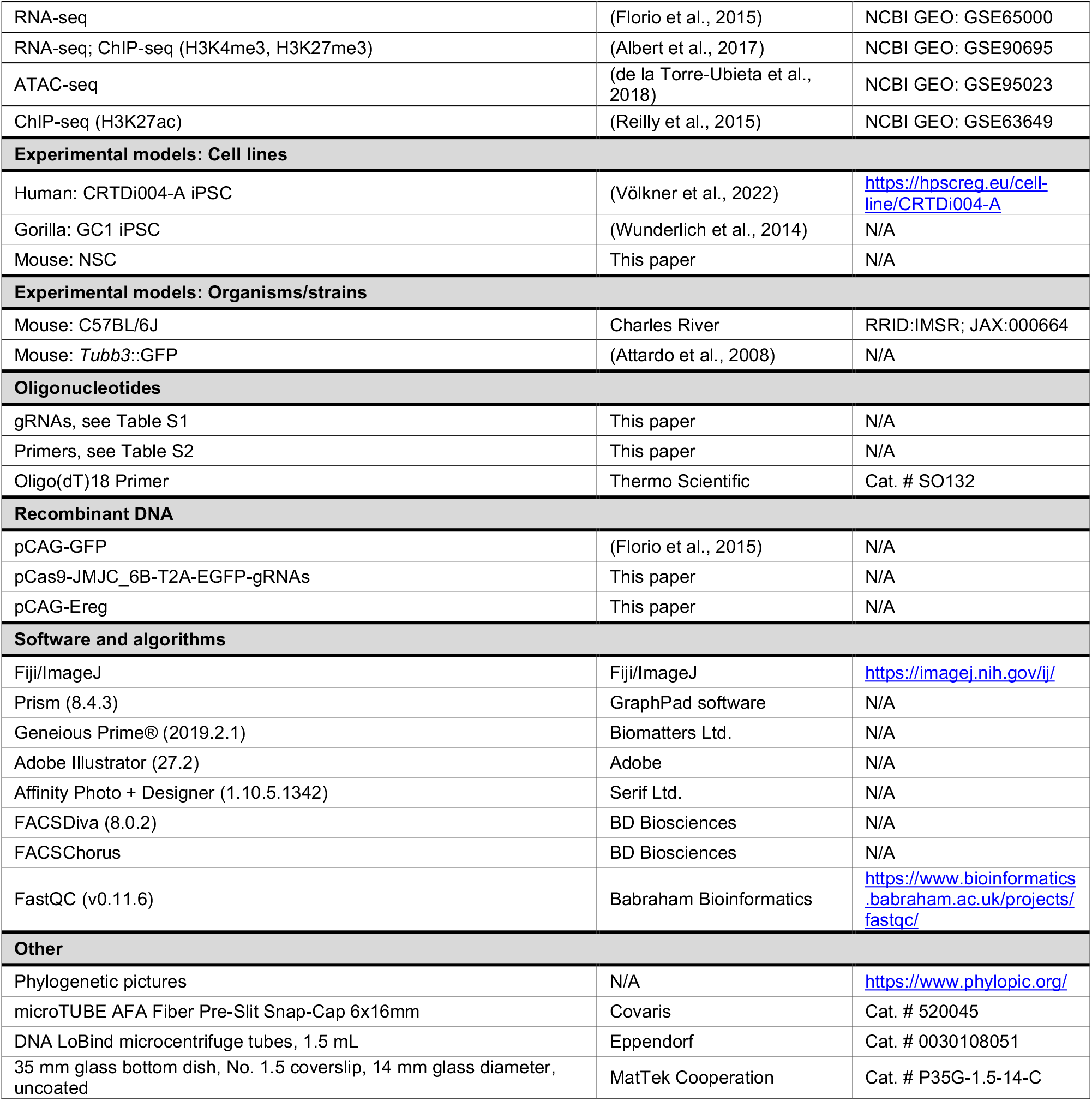

