## Supplemental material for "Primate-expressed EPIREGULIN promotes basal progenitor proliferation in the developing neocortex"

### SUPPLEMENTARY INFORMATION

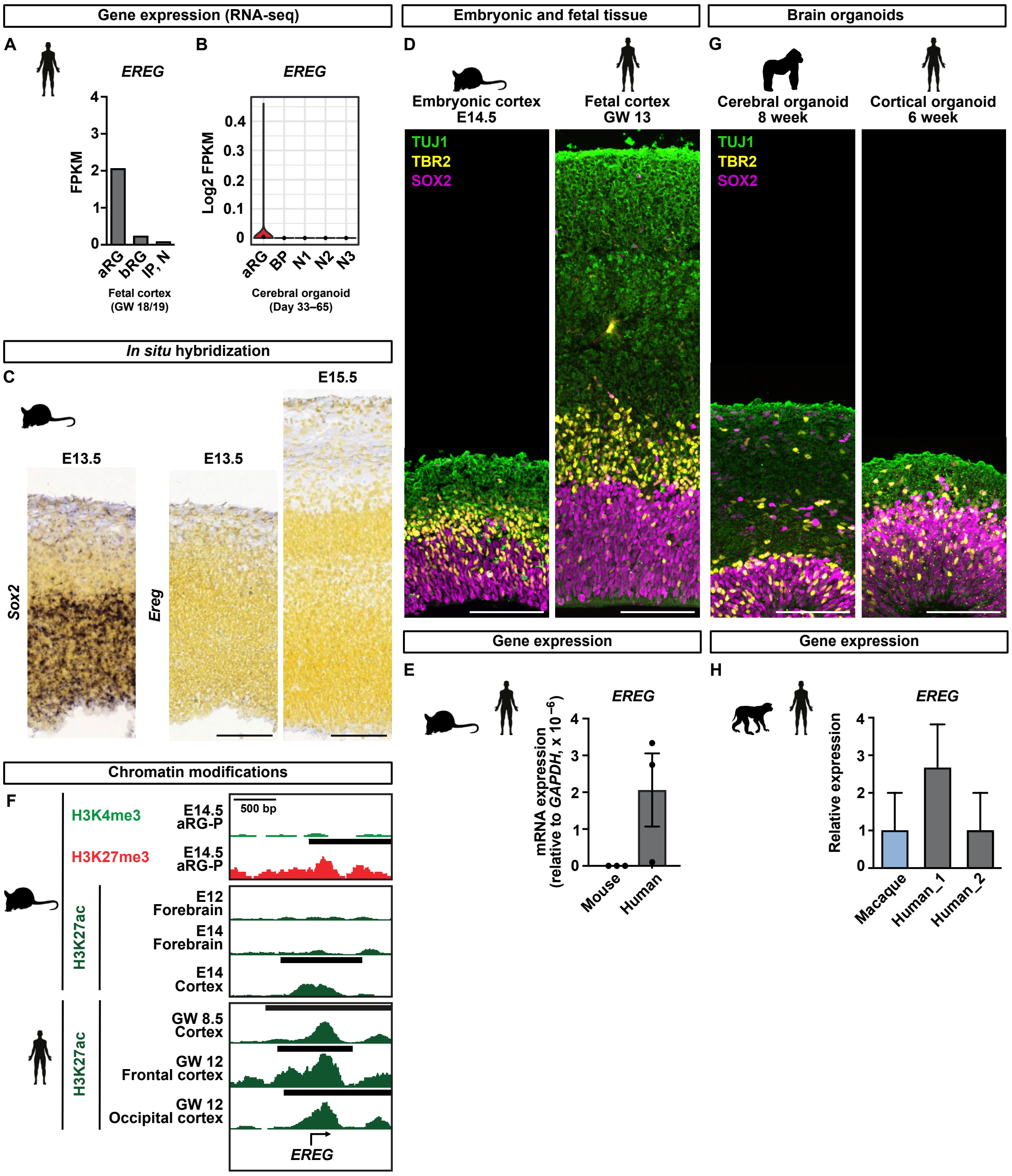


**Supplementary Figure 1 related to Figure 1. Expression of *EREG* in mouse, primate and human neural progenitor cells**

(A, B) *EREG* mRNA levels in human NPCs and neurons from fetal tissue (A) or cerebral organoids (B) analyzed by RNA-seq (data from Camp et al. (2015); Johnson et al. (2015)). (C) *In situ* hybridization data for *Sox2* and *Ereg* of E13.5 and E15.5 mouse neocortex, obtained from the Allen Brain Atlas (Allen Institute for Brain Science, 2004). Scale bars, 100 µm. (D) Immunofluorescence for the RG marker SOX2, the bIP marker TBR2 and the neuronal marker TUJ1 of mNcx and hNcx tissue. Scale bars, 100 µm. (E) RT-qPCR expression analysis of *EREG* in mNcx (E14.5) and fetal hNcx (GW 12/13), relative to *GAPDH*. Error bars represent SD of three biological replicates. (F) H3K4me3, H3K27me3 and H3K27ac ChIP-seq signal around the *EREG* transcription start site (± 1 kb) in mouse proliferative aRG, forebrain and cortex (top) and human cortex (bottom) (data from Albert and Huttner (2018); Gorkin et al. (2020); Reilly et al. (2015). (G) Immunofluorescence for SOX2, TBR2 and TUJ1 of gorilla cerebral and human cortical organoids. Scale bars, 100 µm. (H) *EREG* mRNA levels in macaque and human NPCs analyzed by RNA-seq (data from Kliesmete et al. (2023)).

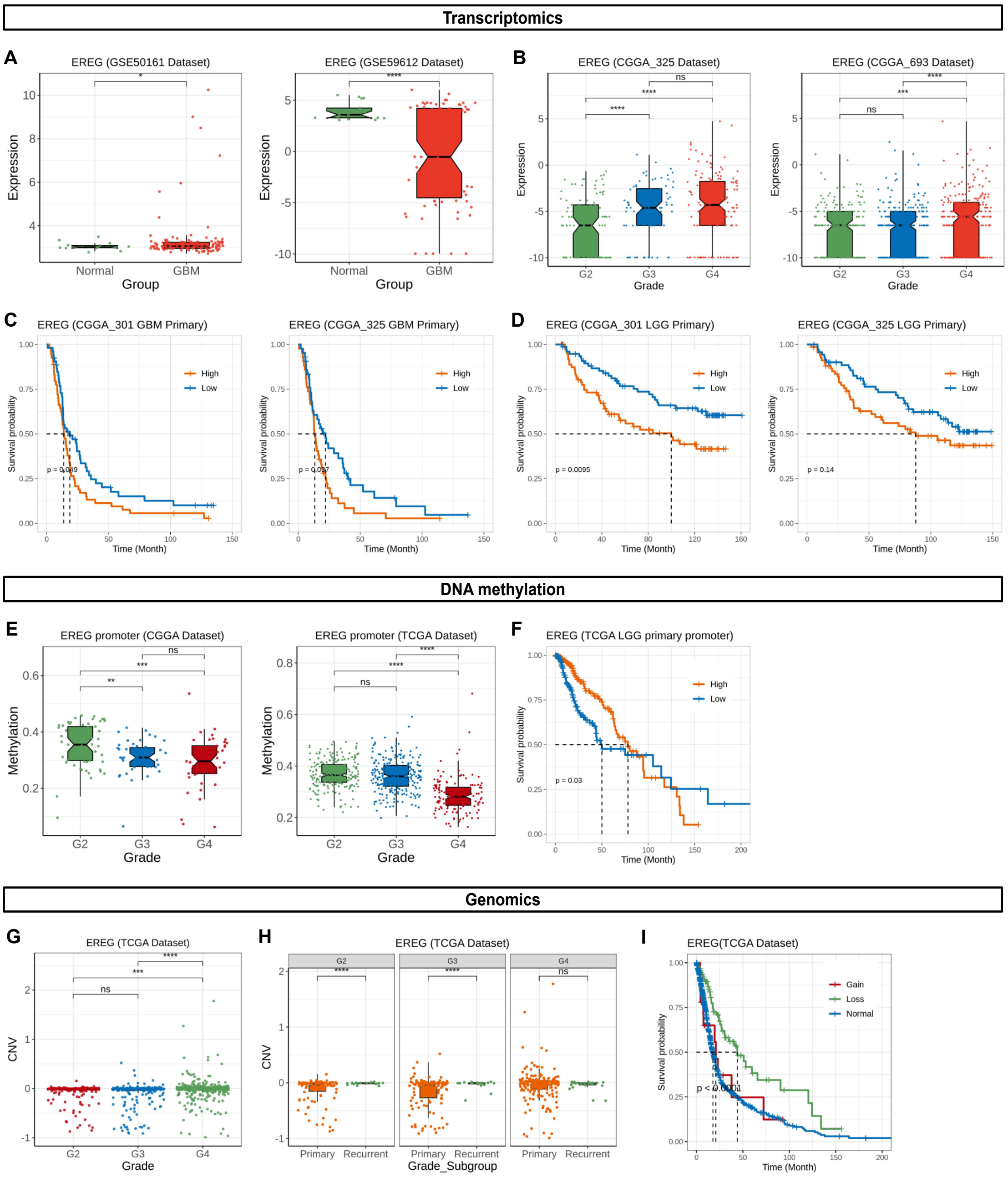


**Supplementary Figure 2 related to Figure 1. Expression, DNA methylation and genomics of *EREG* in brain tumors**

(A) Expression of *EREG* in normal versus glioblastoma (GBM) tissue based on microarray and RNA-seq data (Gill et al., 2014; Griesinger et al., 2013). (B) Expression of *EREG* in grade 2 glioma (G2), G3 and G4 based on RNA-seq data (Bao et al., 2014). (C, D) Survival probability in relation to low (blue) and high (orange) *EREG* expression detected by microarray (Sun et al., 2014; Yan et al., 2012) or RNA-seq (Bao et al., 2014) upon glioblastoma multiforme (C; GBM) and primary low-grade glioma (D; LGG). (E) Promoter DNA methylation of *EREG* in G2, G3 and G4 glioma based on DNA methylation profiling (Ceccarelli et al., 2016; Zhang et al., 2013). (F) Survival probability in relation to low (blue) and high (orange) *EREG* promoter DNA methylation primary low-grade glioma (Ceccarelli et al., 2016). (G, H) Copy number variation (CNV) associated with the *EREG* locus in G2, G3 and G4 glioma (G) and primary versus recurrent glioma (H) (Ceccarelli et al., 2016). (I) Survival probability in relation to gain (red), normal (blue) and loss (green) of *EREG* (Ceccarelli et al., 2016). All data was extracted from BrainBase (https://ngdc.cncb.ac.cn/brainbase/).

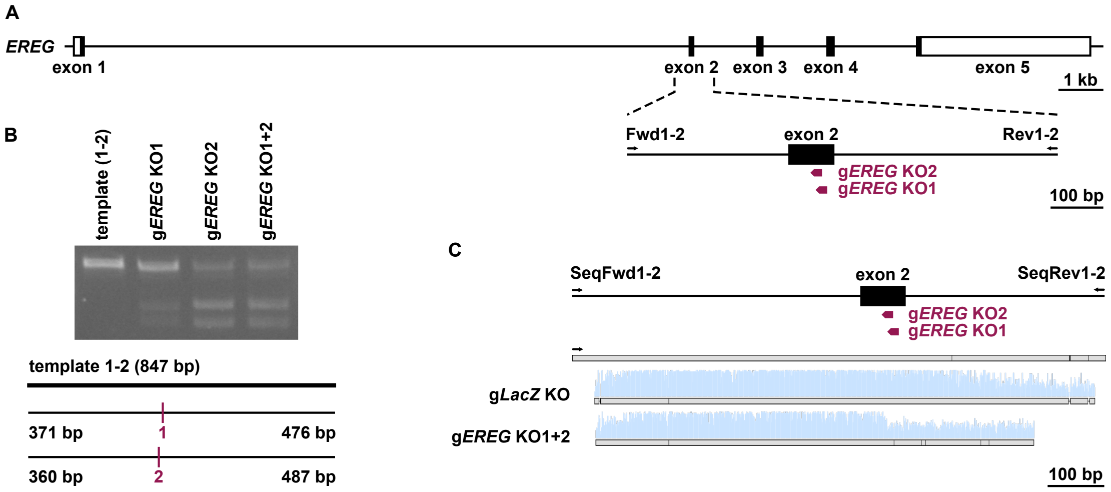


**Supplementary Figure 3 related to Figure 3. Validation of *EREG* gRNA function**

(A) Schematic illustration of the human *EREG* gene locus. The location of the guide RNAs for CRISPR/Cas9-mediated ablation of EPIREGULIN expression (g*EREG* KO1+2) is shown as well as the location of primer binding sites (Fwd, forward; Rev, reverse) for the generation of DNA templates for *in vitro* gRNA efficiency testing. (B) Guide RNA efficiencies were tested *in vitro*. The effects of the g*EREG* KO1+2 RNAs to direct Cas9-mediated cutting of PCR templates was analyzed by agarose gel electrophoresis. Schemes of the sizes of PCR templates, guide RNA binding sites and expected sizes of cut fragments are indicated below. (C) CRISPR/Cas9-mediated targeting of *EREG* was confirmed in iPSCs by electroporation of Cas9/gRNA ribonucleoprotein complexes together with a GFP plasmid, followed by FACS of GFP-positive cells, PCR amplification of the target region and Sanger sequencing. The sequencing results are shown for g*EREG* KO1+2.

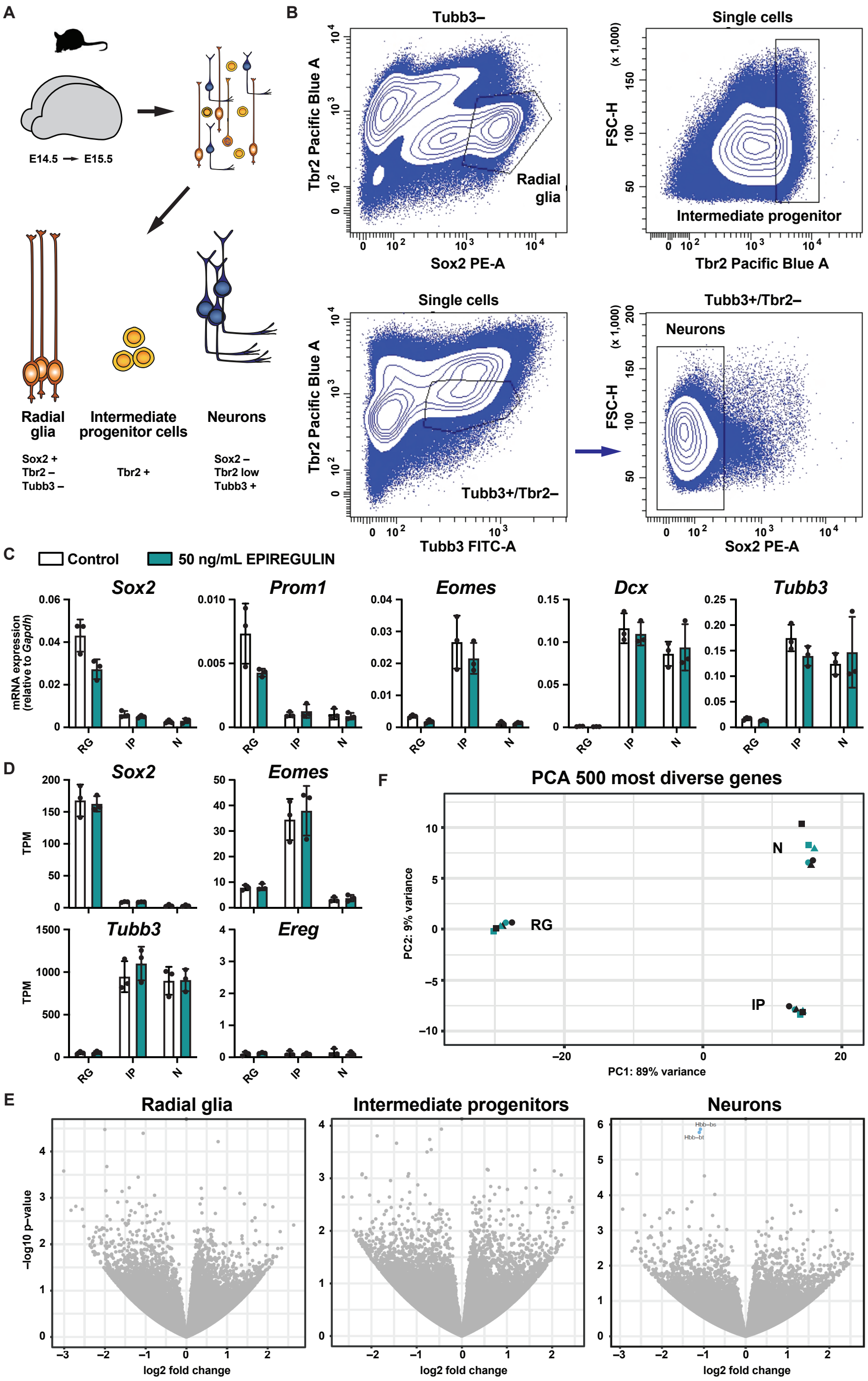


**Supplementary Figure 4 related to Figure 6. Gene expression analysis upon addition of EPIREGULIN to the mouse neocortex.**

(A) Schematic illustration of the experimental workflow. Mouse brain hemispheres (E14.5) from the *Tubb3*::GPF line (Attardo et al., 2008) were isolated and cultured under rotation in the presence of 50 ng/mL of EPIREGULIN for 24 h, dissociated, stained for Sox2 and Tbr2, and cell populations isolated by immuno-FACS based on the indicated marker combinations. (B) Gating strategy of RG (top, left) based on high levels of Sox2 and low levels of Tbr2; bIP (top, right) based on high levels of Tbr2, irrespective of other markers; and neurons (bottom) based on enrichment of GFP expressed from the *Tubb3* promoter and low level of Tbr2, followed by exclusion of Sox2–positive cells. (C) Confirmation of cell type identity by RT-qPCR expression analysis of marker genes characteristic of RG (*Sox2*, *Prom1*), IP (*Eomes*) and neurons (*Dcx*, *Tubb3*) for control and hemispheres treated with EPIREGULIN for 24 hours relative to *Gapdh*. Error bars represent SD of three biological replicates. (D) Expression of *Sox2*, *Eomes*, *Tubb3* and *Ereg* in RG, IP and neurons analyzed by RNA-seq. (E) Volcano plots of log10 (p value) against log2 fold change representing the differences in gene expression in the indicated cell types analyzed by RNA-seq. Grey, non-significant; blue, down-regulated. (F) Principal component analysis (PCA) based on the 500 most divergent genes. The percentage of variance covered by the first two components is indicated.

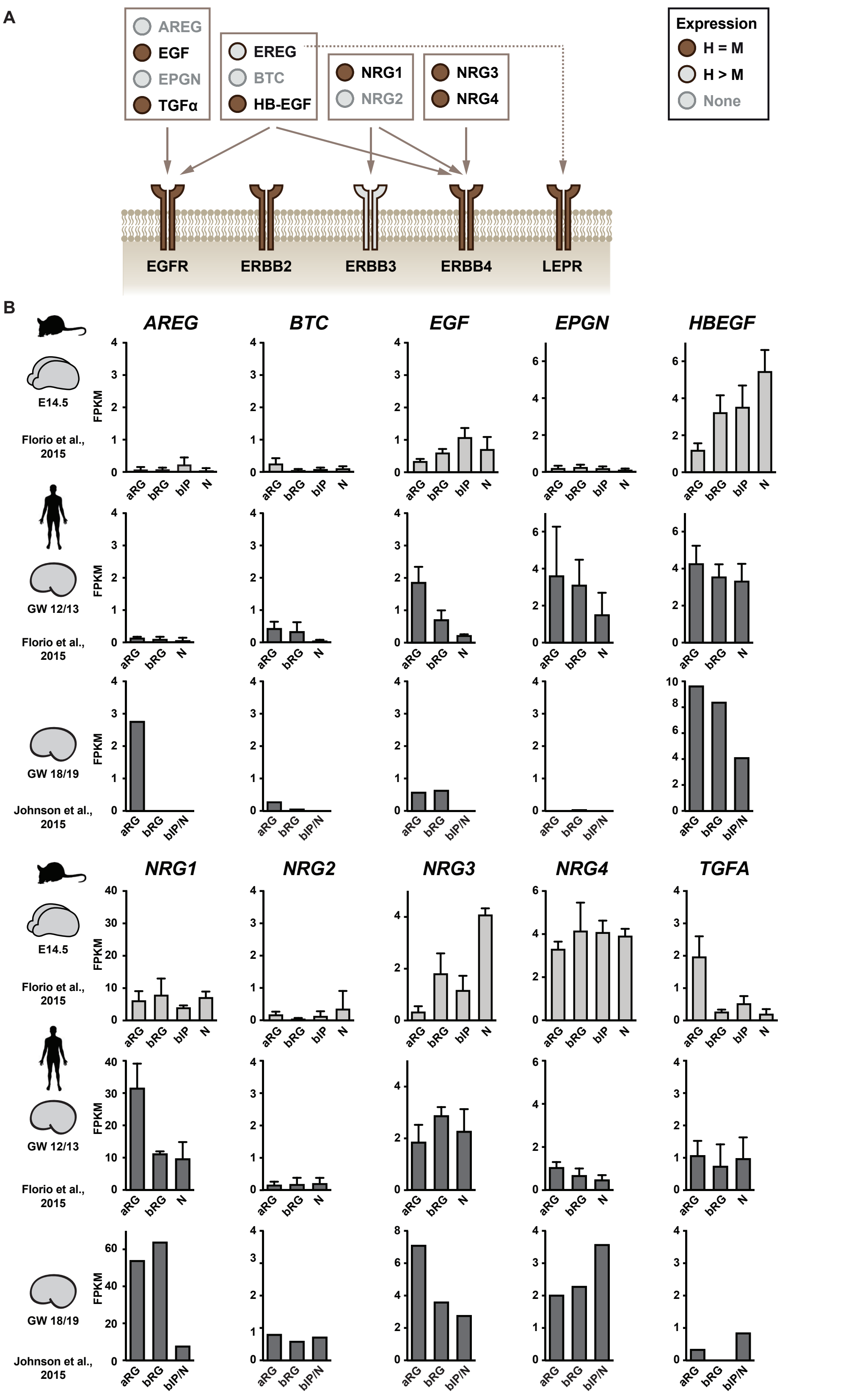


**Supplementary Figure 5 related to Figure 6. Epidermal growth factor and receptor expression**

(A) Schematic illustration of the epidermal growth factors and their respective receptors. Colors indicated mRNA expression: brown, similar in mouse and human; grey with brown contour, higher expression in human RG than in mouse; grey, no expression in both species. (B) Expression of epidermal growth factor genes in mouse (top) and human (bottom) NPCs and neurons analyzed by RNA-seq (data from Florio et al. (2015); Johnson et al. (2015)) at the indicated developmental stages.

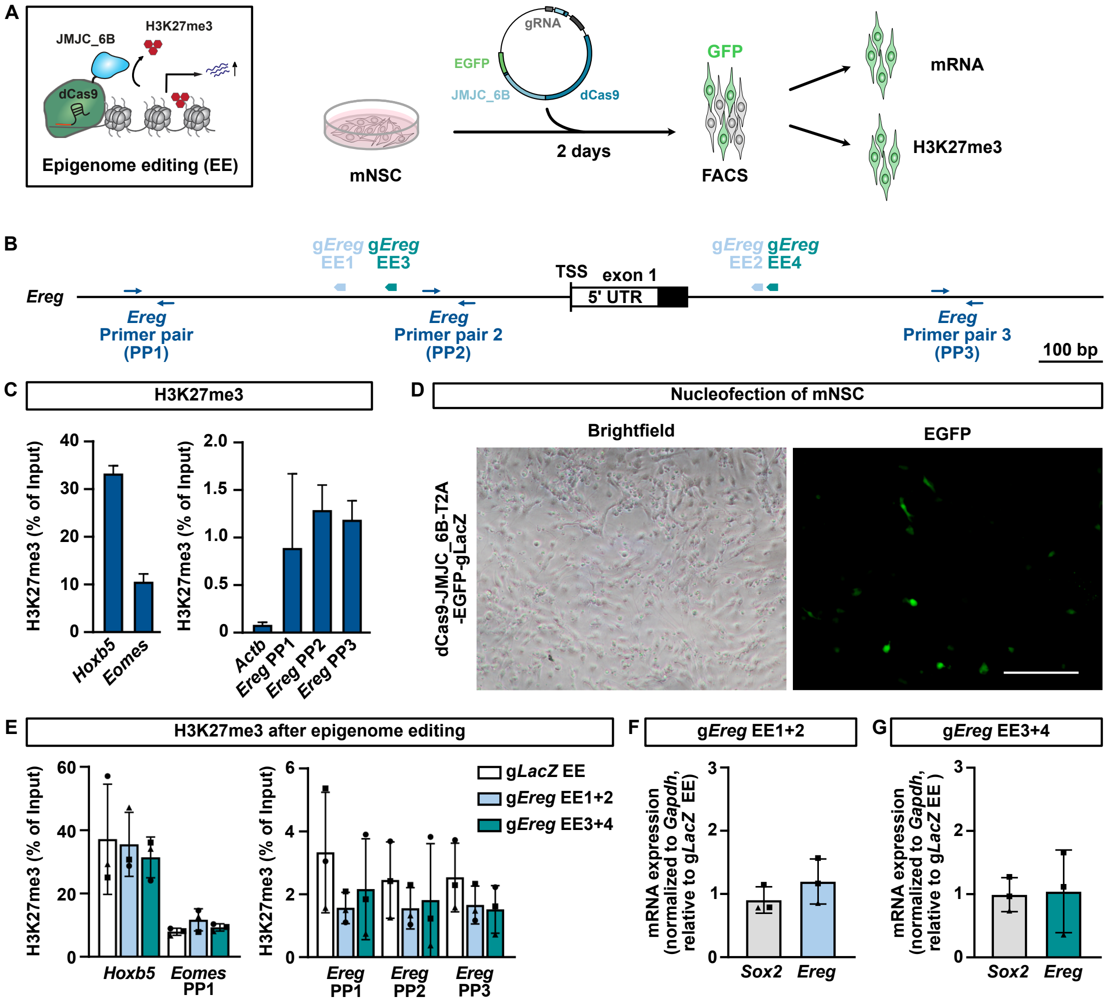


**Supplementary Figure 6 related to Figure 7. Editing of histone methylation at the *Ereg* locus in mNSCs**

(A) Epigenome editing (EE) employing the catalytic domain of KDM6B (JMJC_6B) fused to nuclease deficient Cas9 (dCas9) in mNSCs. Histone methylation and gene expression were analyzed after 2 days following FACS isolation of GFP-positive cells. (B) Location of the gRNAs and primer binding sites (PP, primer pair) for ChIP-qPCR is shown for the *Ereg* locus. Guide RNAs g*Ereg* EE1+2 and g*Ereg* EE3+4 were co-expressed from one plasmid, respectively. (C) Level of H3K27me3 at *Hoxb*, *Eomes, Actb* and *Ereg* (PP1 to PP3) as determined by ChIP-qPCR in mNSCs. (D) Bright field (left) and GFP fluorescence (right) images of mNSCs 2 days following nucleofection with a dCas9-JMJC_6B-T2A-EGFP-gLacZ plasmid. Scale bar, 100 µm. (E) ChIP-qPCR analysis of H3K27me3 around the TSS of *Ereg* and two unrelated genes (*Hoxb5*, *Eomes*) after epigenome editing at the *Ereg* locus. (F, G) Expression of *Sox2* and *Ereg* as determined by RT-qPCR upon epigenome editing using g*Ereg* EE1+2 (F) and g*Ereg* EE3+4 (G). Expression normalized to *Gapdh* and relative to g*LacZ* EE. Error bars represent SD of 3 replicates (from 2-3 independent experiments). One-way ANOVA with Dunnett post hoc test; no statistically significant changes were detected.

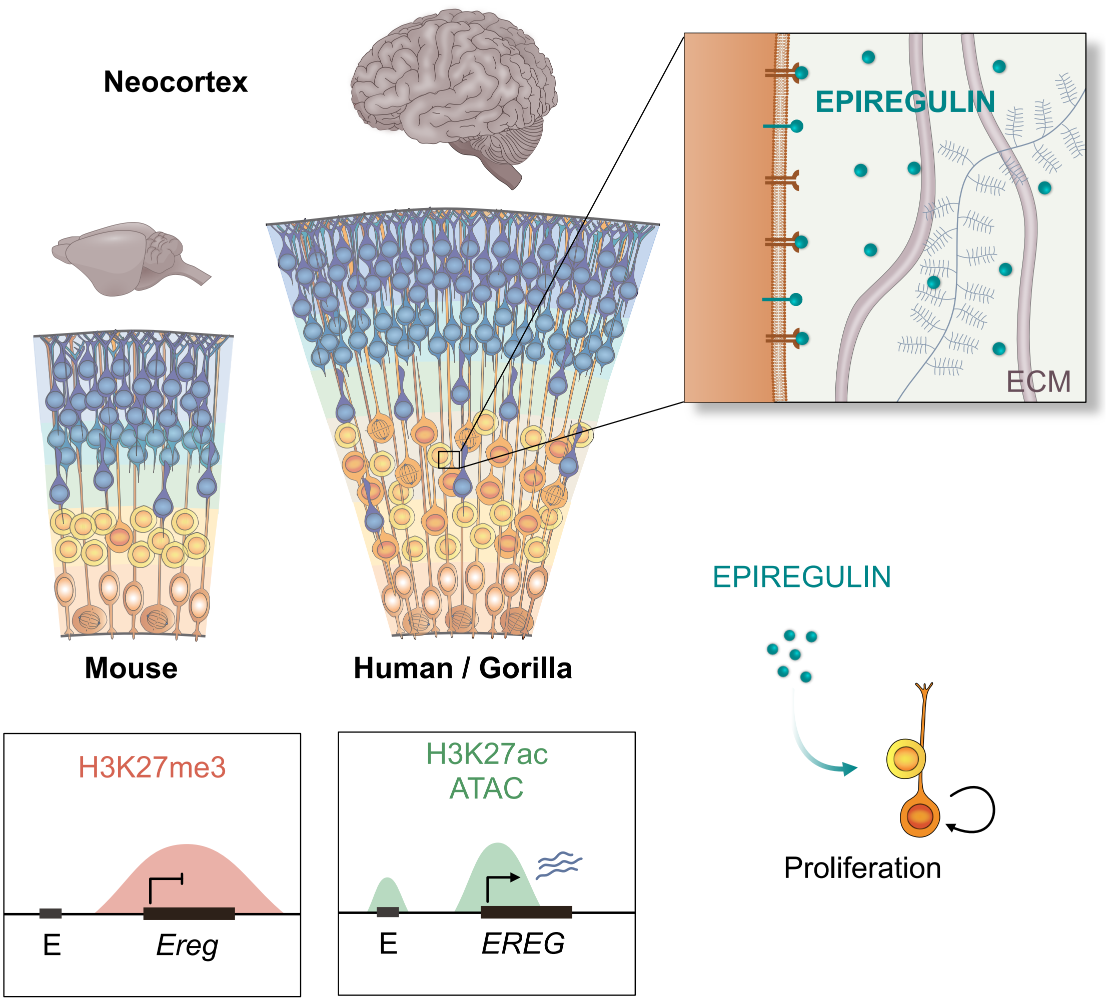


**Supplementary Figure 7 related to Figure 7. Model of EPIREGULIN-mediated regulation of BP proliferation in the neocortex of different species**
Schematic illustration of the expression of *EREG*, encoding the growth factor EPIREGULIN, in the developing mouse, gorilla and human neocortex. While the *Ereg* locus is in a repressive epigenetic state in the mNcx, the human *EREG* gene harbors 11 putative active enhancer regions in its vicinity that may contribute to *EREG* expression in primates. EPIREGULIN induces the proliferation of basal progenitor cells via EGFR-mediated signaling which may represent a mechanism to regulate neocortex size across species.

**Supplementary Table 1. List of gRNAs used.**

| Guide RNA | Sequence | Reference |
|---|---|---|
| Knockout (KO) | | |
| g*LACZ* KO | 5'-TGCGAATACGCCCACGCGATCGG | Kalebic et al, Cell Stem Cell 2019 |
| g*EREG* KO1 | 5'-AGTTATCACTGGACTCTCCTGGG | This paper |
| g*EREG* KO2 | 5'-GACTCTCCTGGGATACATGATGG | This paper |
| Epigenome editing (EE) | | |
| g*LacZ* EE | 5'-TGCGAATACGCCCACGCGAT | Kalebic et al, EMBO 2016 |
| g*Ereg* EE1 | 5'-GCCGGAATAACTCCCTGTTT | This paper |
| g*Ereg* EE2 | 5'-GTCTTTCTCACTGGCGGGGC | This paper |
| g*Ereg* EE3 | 5'-AACTAAAGCCACCCACTACC | This paper |
| g*Ereg* EE4 | 5'-AAGTGCGGGATGGTGCGACT | This paper |

Underlined nucleotides, PAM

**Supplementary Table 2. List of primers used.**

| Oligo | Sequence | Reference |
|---|---|---|
| PCR template for in vitro gRNA efficieny test | | |
| EREG Fwd1-2 | 5'-AAATTCTCCAACTGATGTGC | This paper |
| EREG Rev1-2 | 5'-ACTGTGCCTAGAAAGAAACG | This paper |
| PCR template for sequencing | | |
| **EREG SeqFwd1-2** | 5'-GGATGCCTGTTCCTCCTCATA | This paper |
| EREG SeqRev1-2 | 5'-GAGAGAGAGAGGGGTAGAAAG | This paper |
| RT-qPCR | | |
| Gaphd_F | 5'-TGAAGCAGGCATCTGAGGG | Florio et al., Science 2015 |
| Gaphd_R | 5'-CGAAGGTGGAAGAGTGGGAG | Florio et al., Science 2015 |
| Sox2_F | 5'-TCCCCCCTTTTATTTTCCGTAG | Schmitz et al., EMBO J 2011 |
| Sox2_R | 5'-CCTGATTCCAATAACAGAGCCG | Schmitz et al., EMBO J 2011 |
| Prom1_2F | 5'-CCTCCTGGTGATTTGTCTGC | Florio et al., Science 2015 |
| Prom1_2R | 5'-TTGATCCGAGTCCTGGTCTG | Florio et al., Science 2015 |
| Eomes (Tbr2_2F) | 5'-GACCTCCAGGGACAATCTGA | Florio et al., Science 2015 |
| Eomes (Tbr2_2R) | 5'-GTGACGGCCTACCAAAACAC | Florio et al., Science 2015 |
| Dcx_1F | 5'-TTCAGGACCACAAGCAATGA | Florio et al., Science 2015 |
| Dcx_1R | 5'-GGAAACCGGAGTTGTCAAAA | Florio et al., Science 2015 |
| Tubb3_1F | 5'-ACTTGGAACCTGGAACCATGG | Schmitz et al., EMBO J 2011 |
| Tubb3_1R | 5'-GGCCTGAATAGGTGTCCAAAGG | Schmitz et al., EMBO J 2011 |
| qEreg_1F | 5'-GGCAGTTATCAGCACAACCG | This paper |
| qEreg_1R | 5'-CATCGCAGACCAGTGTAGCC | This paper |
| qEREG_3F_p | 5'-GGTTTCCATCTTCTACAGGCAGT | This paper |
| qEREG_3R_p | 5'-CACCACACGTGGATTGTCTTC | This paper |
| ChIP-qPCR | | |
| Hoxb5_F | 5'-CGACCACGATCCACAAATCAAG | Albert et al., EMBO J 2017 |
| Hoxb5_R | 5'-ATTTGGATAACGCCCCGAGAAG | Albert et al., EMBO J 2017 |
| Eomes_F | 5'-TAAGTCGTGGGTCTGGTCAC | Albert et al., EMBO J 2017 |
| Eomes_R | 5'-CAGCTCTTTCTCCCGTCTGA | Albert et al., EMBO J 2017 |
| Actb_F | 5'-CCCAACACACCTAGCAAATTAGAACCAC | Schmitz et al., EMBO J 2011 |
| Actb_R | 5'-CCTGGATTGAATGGACAGAGAGTCACT | Schmitz et al., EMBO J 2011 |
| Ereg PP1_F | 5'-GCAGAAATGGTGAGGTGGTG | This paper |
| Ereg PP1_R | 5'-CTCTGTGCCTGTCTCAAATGC | This paper |
| Ereg PP2_F | 5'-GGGGCTTGAGTCCGAAGAC | This paper |
| Ereg PP2_R | 5'-GGAACACCTGAGAGGAGGGT | This paper |
| Ereg PP3_F | 5'-CCAGTTTACAACCAGGGGGC | This paper |
| Ereg PP3_R | 5'-TGTTGGCAGTTGCTTCCTGG | This paper |
| RNA-seq library prepartion | | |
| dT-primer | C6-aminolinker-AAGCAGTGGTATCAACGCAGAGTCGACTTTTTTTTTTTTTTTTTTTTTTTTTTTTTTVN | |
| | (where N represents a random base and V any base beside thymidine) | |
| TSO-primer | AAGCAGTGGTATCAACGCAGAGTACATrGrGrG | |
| | (where rG stands for ribo-guanosine) | |
| UP-primer | AAGCAGTGGTATCAACGCAGAGT | |
| Cloning of enhancer candidates | | |
| pMPRA1_BB_F | 5'-TCGGCGGCCAAGCTAGTC | |
| pMPRA1_BB_R | 5'-CAGTTAGGCCAGAGAAATGTTCTGG | |
| RVprimer3(pMPRA1)_F | 5'-CTAGCAAAATAGGCTGTCCC | |
| EBV-rev(pMPRA1)_R | 5'-GTGGTTTGTCCAAACTCATC | |

**Supplementary Table 3. List of CRE sequences tested.**

| Candidate regulatory element | Sequence | Genomic region | | |
|---|---|---|---|---|
| RE_6_human_hg38 | 5'-<br>GTCTCAGATCAAAGAGTCGCCCAAGAATGTGCTTATTAGGAGAACCCACACTCA<br>CCTGGAAACCGGCAGGGGTGGCATCTGGTGACTCAGGCACTGCCCTGCCCACA<br>CCTTCAGTCACTCTCTGCTATAATCCTTGCCAGATCTGTCATTTCAGTGCACAAGC<br>TTGCTTATGTTGCCTCCCAGCTTGAGAACTGACACTGACTCCCCATTGTGCACC | chr4 | 74320682 | 74320898 |
| RE_6_mouse_mm10 | 5'-<br>GTCTCAGATCAAAGAGTCGCCCAAGAATGTGCTTATTAGGAGAACCCACACTCA<br>CCTGGAAACCGGCAGGGGTGGCATCTGGTGACTCAGGCACTGCCCTGCCCACA<br>CCTTCAGTCACTCTCTGCTATAATCCTTGCCAGATCTGTCATTTCAGTGCACAAGC<br>TTGCTTATGTTGCCTCCCAGCTTGAGAACTGACACTGACTCCCCATTGTGCACC | chr5 | 91038606 | 91038822 |
| RE_9_human_hg38 | 5'-<br>AGTTTATTTCACTGTATTCTCCCCAAAAATCTATAAGATAGGAAAAATTAACACCC<br>AGAAAAGTAAATTAATTTTTTTTCAAGATCACGTAGCTCAAGTTTTCCTCACAAGA<br>CTGCCTTTCCACCATAATTTGTGGCCTCTATGTTGGGTTAAATACTCTACTTCAAT<br>TACAGCTATCATAATTTGTTCTCTGTGATGTCCATTATACTCTAGTATT | chr4 | 74377180 | 74377396 |
| RE_9_mouse_mm10 | 5'-<br>AGAACAATCCATTCAGTCTCCTTAAAAGTCTGTCAGAACAATTGAGAGTTGGGG<br>GAGAGAGTTACTGTGTCCAAGTAACAGAGCTCAGGTTTTCTCATGCCATTCTCAT<br>TTCCCCATTCCTTTTGAGGTGATGTAAATTATGCATTAGCTTCTGTAGTGTGTTCC<br>ATGGGATAATCATCACATTCAATATCACTACCCATTTCCAAACTCTGTCCTG | chr5 | 91085587 | 91085803 |
| **Gibson overhangs** | **Sequence** | | | |
| Forward | TGCCAGAACATTTCTCTGGCCTAACTGGCC AGGACCGGATCAACT | | | |
| Reverse | CTGAGCAT TGGA GC GTCTTCGAATTCATCTGGTACC TCGGTTCACGCAATG | | | |
